# Neuroinflammation characterizes the earliest changes in Alzheimer’s disease pathology and associated subjective cognitive impairment in adult hydrocephalus biopsies

**DOI:** 10.1101/2020.10.01.322511

**Authors:** Wenrui Huang, Anne Marie Bartosch, Harrison Xiao, Xena Flowers, Sandra Leskinen, Zeljko Tomljanovic, Gail Iodice, Deborah Boyett, Eleonora Spinazzi, Vilas Menon, Robert A. McGovern, Guy M. McKhann, Andrew F. Teich

**Affiliations:** Columbia University, Department of Pathology and Cell Biology, New York, NY; Columbia University, Taub Institute for research on Alzheimer’s Disease and the Aging Brain, New York, NY; Columbia University, Department of Neurology, New York, NY; Columbia University, Department of Neurosurgery, New York, NY; University of Minnesota, Department of Neurosurgery, Minneapolis, MN

## Abstract

In an effort to better characterize the transcriptomic changes that accompany early Alzheimer’s disease (AD) pathology in living patients and correlate with contemporaneous cognitive data, we performed RNA-seq on 106 cortical biopsies that were taken during shunt placement for adult onset hydrocephalus with varying degrees of comorbid AD pathology. A restricted set of genes correlate with AD pathology in these biopsies, and co-expression network analysis demonstrates an evolution from microglial homeostasis to a disease-associated microglial phenotype in conjunction with increasing AD pathologic burden, along with a subset of additional astrocytic and neuronal genes that accompany these changes. Further analysis demonstrates that these correlations are driven by patients that report mild cognitive symptoms, despite similar levels of β-amyloid and tau pathology in comparison to patients who report no cognitive symptoms. Interestingly, downregulation of homeostatic genes and upregulation of disease-associated genes also correlate with microglial plaque invasion and an activated microglial morphology, and this change is not sensitive to cognitive status, suggesting that an initial microglial response to AD pathology is eventually maladaptive. Taken together, these findings highlight a restricted set of microglial and non-microglial genes and suggest that early AD pathology is largely characterized by a loss of homeostatic genes and an activated microglial phenotype that continues to evolve in conjunction with accumulating AD pathology in the setting of subjective cognitive decline.

## Introduction

Recent interest in RNA-seq has led to the generation of several large datasets of transcriptomic data from autopsy Alzheimer’s disease (AD) brain tissue ^1, 2, 3, 4^. Analysis of this data has produced a wealth of information about a wide range of pathophysiologic processes in AD. The goal of the present study is to add to this data by exploring the changes in gene expression accompanying early-stage AD pathology in surgical biopsies without post-mortem artifact, and compare to contemporaneously gathered cognitive data. To do this, we used biopsies taken from patients during surgical shunt placement to treat chronic hydrocephalus at the Columbia University adult hydrocephalus clinic. Chronic hydrocephalus in the aging population can occur for a variety of reasons, although the etiology is often unclear. In the absence of a clear etiology, most of these cases are categorized as “Idiopathic Normal Pressure Hydrocephalus” (or iNPH; see Methods for a further discussion of our NPH cohort). Placing a ventricular shunt is often effective for symptom relief in the setting of NPH/chronic hydrocephalus ^5, 6, 7^, although which patients will have persistent long-term clinical benefit remains to be determined ^8^. At the time of shunt placement, a cortical biopsy is routinely obtained at our center to look for possible coexistent brain pathology. The biopsy is taken at the brain entry point of the ventricular shunt catheter. Perhaps not surprisingly, cortical biopsies taken from elderly NPH patients at shunt placement have been shown to have a relatively high frequency of β-amyloid plaque pathology, ranging from 42% to 67% ^9, 10^, perhaps due to the fact that early-stage AD in many cases may actually be causing some of the symptoms attributed to NPH/chronic hydrocephalus.

Consistent with this hypothesis, the presence of either 1) severe β-amyloid plaque pathology or a CSF AD signature of high phospho-tau/β-amyloid ratio have both been shown to predict a lack of response to shunting ^10, 11^. Interestingly, unlike β-amyloid plaque pathology, tau pathology is relatively sparse in NPH cortical biopsies ^10^, although some studies have found trace tau pathology at higher levels ^12^, which is consistent with the fact that most patients coming to shunt surgery are not severely demented (those patients that are pre-AD are likely to be at a Braak stage with sparse to no neocortical tangles).

In summary, although several large autopsy cohorts of AD brain data are now publicly available, surgical biopsies from hydrocephalus patients yield several advantages. First, this cohort represents a large sample of relatively young patients, many of which have early stage AD pathology ^9, 10^. Specifically, our cohort has an average age (74.9) that is significantly lower than the average age of existing AD autopsy cohorts, which are typically in the mid to upper 80’s ^1, 4^, and is just under the average age of initial clinical presentation for AD (75.5) ^13^. The tissue in this cohort is also free of gene expression changes that accompany end-of-life hypoxia/agonal state, as well as any changes in RNA caused by post-mortem degradation, and all cognitive data curated from the patients’ charts represents the cognitive state relatively close to the time of tissue acquisition (average time between cognitive exam and biopsy in our cohort is 120 days; see Methods). Despite these unique advantages, it should also be noted that the confounding factor of “hydrocephalus” is present throughout this cohort. Crucially, we validate the findings in this study in AD autopsy cohorts, and further note that comobidity is a common problem in AD autopsy cohorts as well (see Discussion section for further discussion).

In contrast to the large-scale changes in gene expression seen in AD autopsy studies ^1, 2, 3, 4^, analysis of these biopsies shows a restricted set of microglial (and some non-microglial) genes that correlate with AD pathology primarily in patients that report subjective cognitive impairment. In contrast to the existing human AD autopsy literature, we also identify a homeostatic microglial module that replicates the decrease in homeostatic genes that is seen in the mouse AD literature. Finally, these microglial modules also correlate in a coherent way with plaque infiltration and changes in microglial morphology, and this change is not sensitive to cognitive status. Taken together, these data suggest that an initial microglial response to AD pathology is eventually maladaptive, and is associated with accumulating pathology, non-microglial cell responses, and cognitive dysfunction.

## Results

In this study, we examine changes in gene expression that accompany early AD pathology in cortical biopsies that were removed in the course of shunt placement for NPH, and compare the results with AD pathology on histology and contemporaneously gathered cognitive data (see Figure 1 for our workflow and Methods for further details on our cohort). Specifically, we performed RNA-seq on 106 biopsies from NPH patients, with an average age of 74.9. In all cases, patients were shunted for chronic hydrocephalus by the same surgical team, and biopsies were taken from a specific area of frontal cortex (2/3 of our cohort) or parietal cortex (1/3 of our cohort). The decision to shunt/biopsy in frontal or parietal cortex was made by the surgeon based on cosmetic considerations (see Methods). Changes in gene expression that correlate with AD pathology in our samples trend similarly in frontal and parietal cortex (see Supplemental Table 1), and even when we combine all samples, very few individual genes reach statistical threshold (see below).

**Figure 1:**
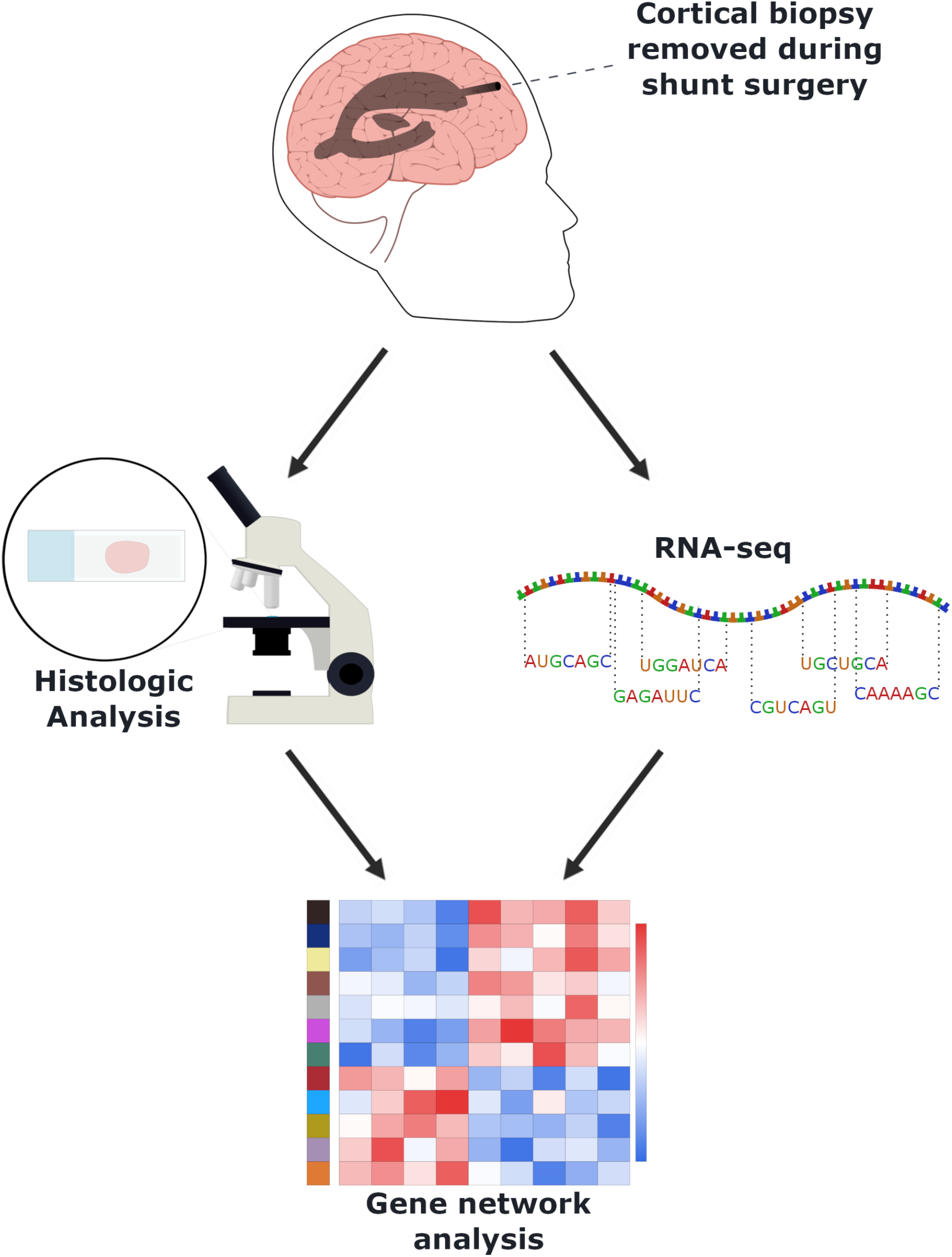
Biopsies removed for ventricular shunting in the operating room are immediately split in half. Half of the biopsy is frozen in liquid nitrogen and sent for RNA-seq. The other half is formalin fixed and paraffin embedded for subsequent pathologic analysis (see Methods).

### Immune/Microglia specific genes are strongly correlated with NPH AD pathology

We initially processed our RNA-seq data by regressing out variability in gene expression not associated with our primary variables of interest (β-amyloid plaque and tau pathology) ^14^. β-amyloid plaques were counted per square millimeter area on slides of tissue immunostained with 6E10 antibody. In order to quantify tau, we devised a rating scale to grade the minimal degree of tau pathology seen in NPH biopsy slides immunostained with AT8 antibody (see Supplemental Figure 1 for examples of each grade). Grade 0 was given to biopsies with no tau pathology (n = 42). Grade 1 was given to biopsies that have any tau pathology at all, usually one or more dystrophic neurites (n = 39). Grade 2 was given to biopsies that have at least one tau-positive neuron or neuritic plaque (n = 18). Grade 3 was reserved for biopsies with tau pathology evenly distributed throughout the biopsy (n = 7).

Initial analysis of the data identified a restricted set of 38 genes that passed FDR of 0.1 at the individual gene level that correlated with either β-amyloid and/or tau burden. Indeed, one of the most striking things about our initial analysis is how much isn’t changing in these biopsies, especially given the large-scale changes in gene expression seen in many other autopsy-based cohorts that include brains with clinical AD ^1, 2, 3, 4^. The genes that pass FDR threshold are enriched for immune response genes, many of which have been tied to AD. Figure 2 shows the top 20 genes that correlate with β-amyloid plaques and tau burden. A number of immune or microglia specific genes were among the Top 20 list, including TREM2 and C4B, both of which have been implicated in the immune response in AD ^15, 16^. These results point to microglia/immune response changes as being important in the very earliest stages of AD, and occur before other physiologic changes appear at the bulk RNA-seq level. A Fisher’s exact test confirmed that microglia-specific genes are overrepresented amongst the genes that individually past our FDR threshold, both using mouse cell-type specific RNA-seq data ^17^ and human single-nucleus RNA-seq data ^18^ (Figure 2, panels C and D and Supplemental Table 1; note that while the trend is similar in both analyses, only the human comparison passes Bonferroni adjustment).

**Figure 2:**
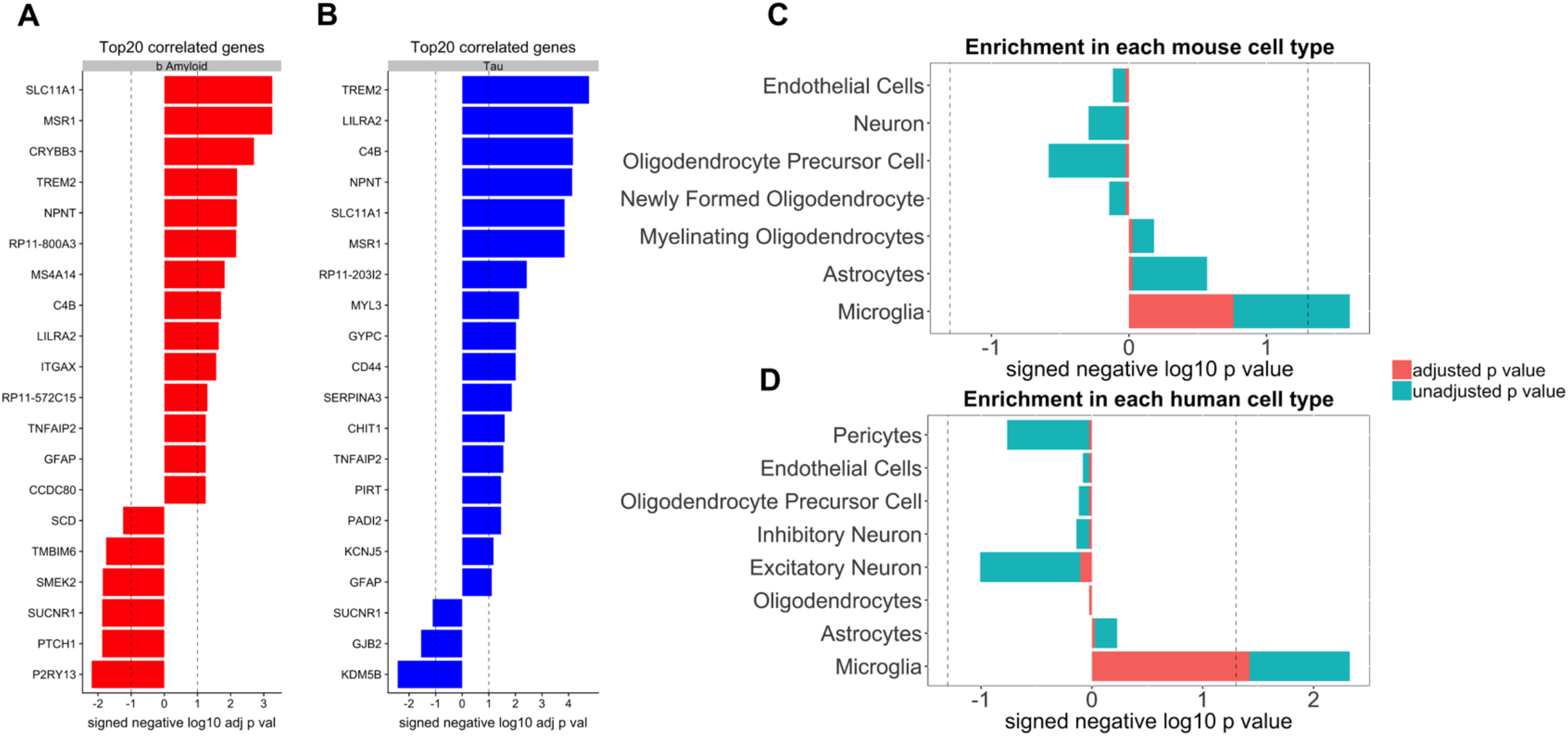
A restricted set of genes correlates with β-amyloid (**Panel A**) and tau (**Panel B**) burden in NPH biopsies (FDR adjusted using the Benjamini-Hochberg procedure across all genes in the transcriptome – dotted line is FDR = 0.1; see Supplemental Table 1 for the full list). **Panels C and D**: A Fisher’s exact test confirmed that microglia-specific genes are overrepresented amongst the genes that individually past our FDR threshold, both using mouse cell-type specific RNA-seq data ^17^ (panel C) and human single-nucleus RNA-seq data ^18^ (panel D; panels C and D show unadjusted and Bonferroni adjusted p-value; dotted line = p-value of 0.05; See Supplemental Table 1 for numerical values; note that while the trend is similar in both analyses, only the human comparison passes Bonferroni adjustment).

### AD pathology gene correlations are strongest in patients with subjective cognitive impairment

Previous work with AD RNA-seq tissue has used gene network analysis to further clarify how groups of genes correlate with various AD traits ^1^. When we performed Weighted Gene Co-expression Network Analysis (WGCNA) ^19^ on our NPH data, we identified in total 58 gene co-expression modules, only three of which are significantly correlated with either β-amyloid or tau burden (*saddlebrown, orange*, and *darkgrey* modules – Figure 3; see Supplemental Table 2 for full results of WGCNA analysis), consistent with analysis at the single-gene level that a restricted set of genes correlate with AD pathologic traits in this cohort. To further refine our analysis, patient charts were curated for data that would help differentiate the patients by cognitive status. Although rigorous cognitive testing was not consistently carried out, the majority of patients and their families were asked whether they had experienced subjective cognitive impairment during an exam close in time to the biopsy date (see Methods). Using this simple metric (yes vs. no), we were able to assign 93 of our sequenced biopsies into these two groups, with 59 replying “yes” and 34 replying “no” (the remaining 13 biopsies came from patients where we were unable to locate the answer to this question in the chart). Patients that reported subjective cognitive impairment had non-significantly higher β-amyloid and tau load than patients that reported no cognitive impairment (p-value for β-amyloid = 0.21; p-value for tau = 0.66 by Mann Whitney U test). Interestingly, all of the tau grade 3 biopsies with cognitive information have a history of subjective cognitive complaint. However, these biopsies are so few in number (7) that they do not significantly affect the overall analysis. Interestingly, there is significantly less co-occurrence of β-amyloid pathology and tau pathology in the same biopsy from patients that report no cognitive impairment when compared to patients that do report subjective cognitive impairment (see Supplemental Figure 2). This suggests that AD pathology may be more widespread in patients that report cognitive impairment (so that we are more likely to see both β-amyloid and tau in a small biopsy), even if the local density of AD pathology in these biopsies is not significantly higher compared to patients that report no cognitive impairment.

**Figure 3:**
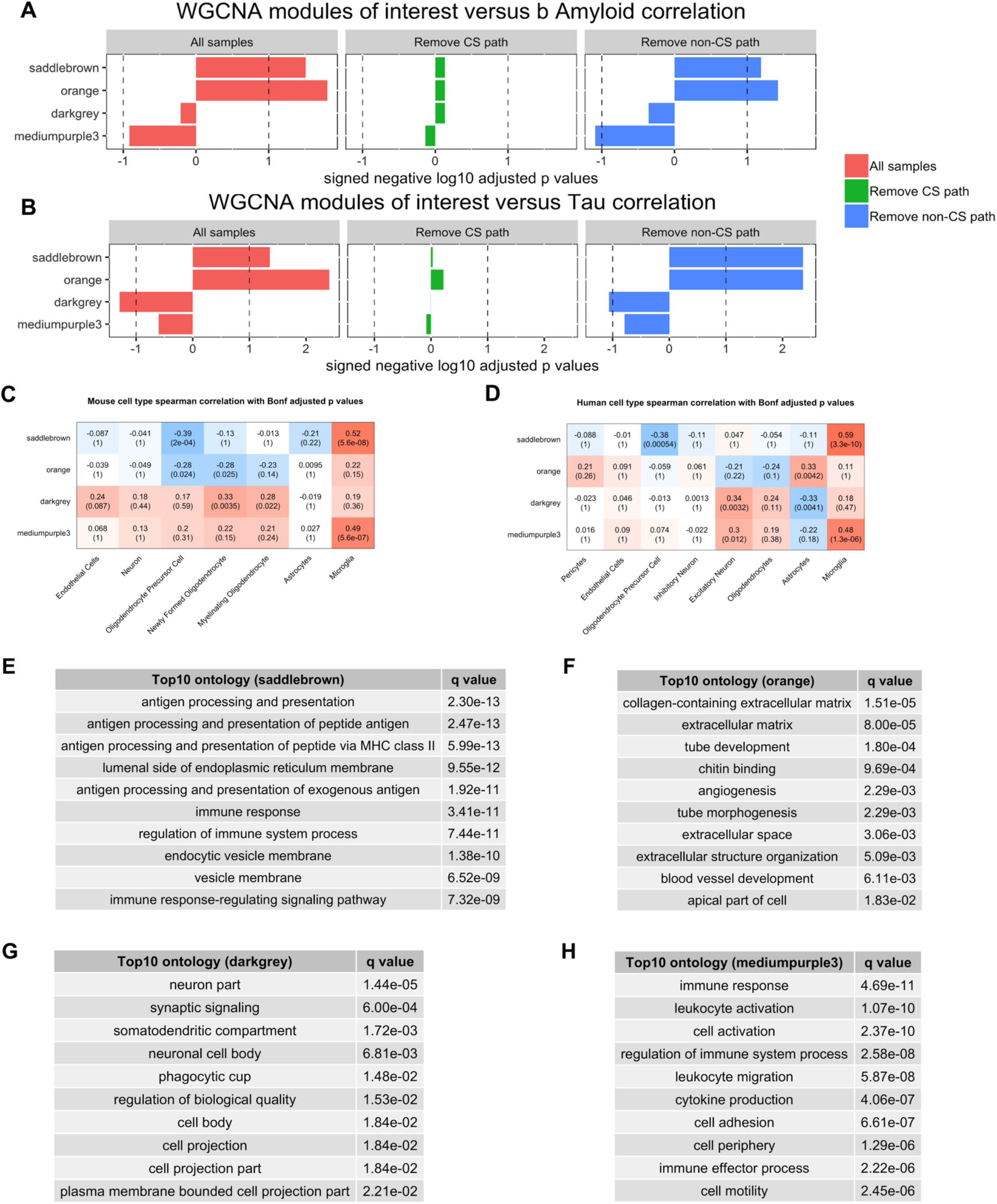
Gene expression modules correlate with AD pathology in the setting of subjective cognitive impairment. **Panels A and B**: Consistent with the single gene analysis, WGCNA shows that only three modules (*saddlebrown, orange, and darkgrey*) correlate significantly with β-amyloid (panel A) and tau (panel B) when all samples are considered (red bars on left; see Supplemental Table 2 for the correlations for all WGCNA modules from this analysis and Supplemental Table 3 for the list of genes in key modules in this paper). Subjective cognitive status strongly influences how microglial modules correlate with β-amyloid and tau pathology (green and blue bars). When we removed all samples with AD pathology from our cohort that reported cognitive impairment, this abolished the significant correlation of the modules with AD pathology (green bars). In contrast, when we do the converse (i.e. remove all samples with AD pathology from our cohort that do not report cognitive impairment), our gene expression modules continue to correlate with AD pathology in the remaining biopsies (blue bars). In fact, one module that was previously not significant becomes significant (*mediumpurple3*). Panels A and B are FDR adjusted using the Benjamini-Hochberg procedure across all 58 modules in our WGCNA analysis - dotted line in A and B is FDR = 0.1. **Panels C and D**: The modules in panels A and B were correlated with cell-type specific gene lists (using mouse cell-type specific RNA-seq data ^17^ (panel C) and human single-nucleus RNA-seq data ^18^ (panel D)). While *saddlebrown* and *mediumpurple3* are clearly microglial, *darkgrey* and *orange* are more weakly neuronal and astrocytic respectively by correlation (each row in panels C and D is separately Bonferroni adjusted; see Supplemental Figure 3 for enrichment of these modules with cell-type specific gene lists, and Supplemental Table 5 and 6 for all correlation and enrichment values and p-values for these modules with mouse and human cell types). **Panels E-H**: The top 10 ontology groups for each module are displayed (see Supplemental Table 7 for full ontology analysis for these four modules).

Although cognitive status does not predict significantly more AD pathology in our cohort, we did find that the correlations of our modules with AD pathology are being driven by patients that report cognitive impairment. When we removed all samples with AD pathology from our cohort that reported cognitive impairment, this abolished the significant correlation of the modules with AD pathology in the remaining biopsies (Figure 3, panels A and B; see Supplemental Table 2 for full analysis with all WGCNA modules). In contrast, when we do the converse (i.e. remove samples with AD pathology that do not report cognitive impairment), our gene expression modules continued to correlate with AD pathology in the remaining biopsies. In fact, an additional module reached the significance threshold (*mediumpurple3*; see Supplemental Table 3 for genes in the modules of interest in this manuscript). To further examine the sensitivity of gene correlations with AD pathology to cognitive status, we ran 1000 iterations where we randomly replaced half of the samples with AD pathology in the analysis shown in blue (Figure 3 panels A and B) with pathology samples in the analysis shown in green (i.e. pathology samples with subjective cognitive impairment are being randomly replaced with pathology samples without documented cognitive impairment). As noted in Supplemental Table 4, this did not statistically change the overall distribution of the burden of pathology in any of the simulations.

In contrast, all four of our modules fail to pass 0.1 FDR significance in their correlation with β-amyloid and tau for the majority of the simulations. Taken together, these findings indicate that the correlations of these modules with AD pathology are highly sensitive to cognitive status.

In an effort to better characterize these modules, we first determined whether these modules correlate with cell-type specific gene lists. Analysis with gene lists from mouse cell-type specific RNA-seq data ^17^ and human single-nucleus RNA-seq data ^18^ both show similar cell class assignments, and this is further supported by enrichment analysis (Supplemental Figure 3 and Supplemental Tables 5 and 6). These analyses support the view that the *saddlebrown* and *mediumpurple3* modules are predominantly microglial. While the data is somewhat more mixed for *darkgrey* and *orange*, the overall trend is that *darkgrey* is neuronal while *orange* is astrocytic, which is broadly consistent with the positive correlation of the *orange* module and negative correlation of the *darkgrey* module with β-amyloid and tau in the NPH data.Ontology analysis is consistent with these observations (Figure 3, panels E-H; see Supplemental Table 7 for full ontology analysis results), with *saddlebrown* and *mediumpurple3* characterized by immune response ontology groups, and *darkgrey* by neuronal groups (note that the *orange* module’s cell-type specificity is less clear from ontology analysis).

In our implementation of WGCNA, we allowed for both positive and negative gene correlations within modules (“unsigned” implementation; see Methods). This allowed us to potentially capture more complex changes in physiology within the same module, rather than only model up or down-regulation of groups of genes together. In addition, using unsigned modules also eliminates the extra analysis step needed to pair signed modules that are anti-correlated with each other. Of the four modules that pass FDR threshold in Figure 3, *darkgrey* is notable for having the largest fraction of genes that negatively correlate with the PC1 eigengene (37% of *darkgrey* genes correlate negatively with the PC1 eigengene, while the other three modules all have less than 20% of their genes negatively correlating with PC1; see Supplemental Table 3). For all four of our modules of interest, ontology analysis of positively correlating genes replicated the groups seen in ontology analysis of the full module, consistent with the positively correlating genes dominating the signature of these modules (see Supplemental Table 7 for all ontology analysis results). On the other hand, ontology analysis of negatively correlating genes revealed no significant groups for *saddlebrown*, one group for *mediumpurple3*, and several groups without a clear theme for *orange*. In contrast, ontology analysis for negatively correlating genes in *darkgrey* revealed several significant groups related to lipid metabolism (including “lipid binding” and “lipid transporter activity”), This suggests that upregulation of lipid metabolism may be an early change that occurs in tandem with early neuronal dysfunction and loss of synaptic/neuronal genes in AD. Lipid metabolism is increasingly recognized as playing an important role in AD pathogenesis ^20, 21^, and two of the genes from these ontology groups (ApoB and PCTP) have recently been implicated in AD though analysis of GWAS data ^22, 23^. In addition to these changes, we also note several other compensatory genes in *darkgrey* that negatively correlate with the PC1 eigengene, such as HSB1 and neuroglobin, which have both been shown to increase in AD and are thought to be part of the stress response ^24, 25, 26^. In summary, the *darkgrey* module suggests that we may be capturing early neuronal changes along with compensatory/reactive changes in these biopsies that correlate with increasing pathology most significantly in patients with subjective cognitive impairment. Moreover, this analysis suggests that while at the single gene level the changes we are observing are overwhelmingly microglial (Figure 2), WGCNA is identifying correlating genes from other cell types that may be less significant at the single gene level.

### Microglial gene expression modules are enriched for previously identified sets of homeostatic and disease-associated immune response genes

In Figure 3, one of our microglial modules (*saddlebrown*) positively correlates with AD pathology, whereas the other microglial module (*mediumpurple3*) negatively correlates with AD pathology. In an effort to characterize these modules further and explain these associations, we determined whether these modules were associated with known sets of disease-associated microglial genes. Specifically, two recent studies (Keren-Shaul et al ^27^ and Mathys et al ^28^) have identified a transition from homeostasis to a late stage/disease-associated microglial phenotype in AD mouse models, suggesting that as AD pathology accumulates, microglia undergo a shift in their transcriptomic profile. Interestingly, *mediumpurple3* strongly correlates with homeostatic gene lists (Figure 4, panel A). In contrast, *saddlebrown* strongly correlates with lists of late-stage/disease-associated microglial genes (Figure 4, panel B; see Supplemental Table 8 for correlation and p-values for all analyses in Figure 4). In addition, *mediumpurple3* is significantly enriched for homeostatic genes, while *saddlebrown* is significantly enriched for DAM stage 2/late response genes (Figure 5 E and G and Supplemental Table 11; note that individual homeostatic and DAM stage 2/late response genes positively correlate with the PC1 eigengene of *mediumpurple3* and *saddlebrown* respectively – see Supplemental Table 14 for details).

**Figure 4:**
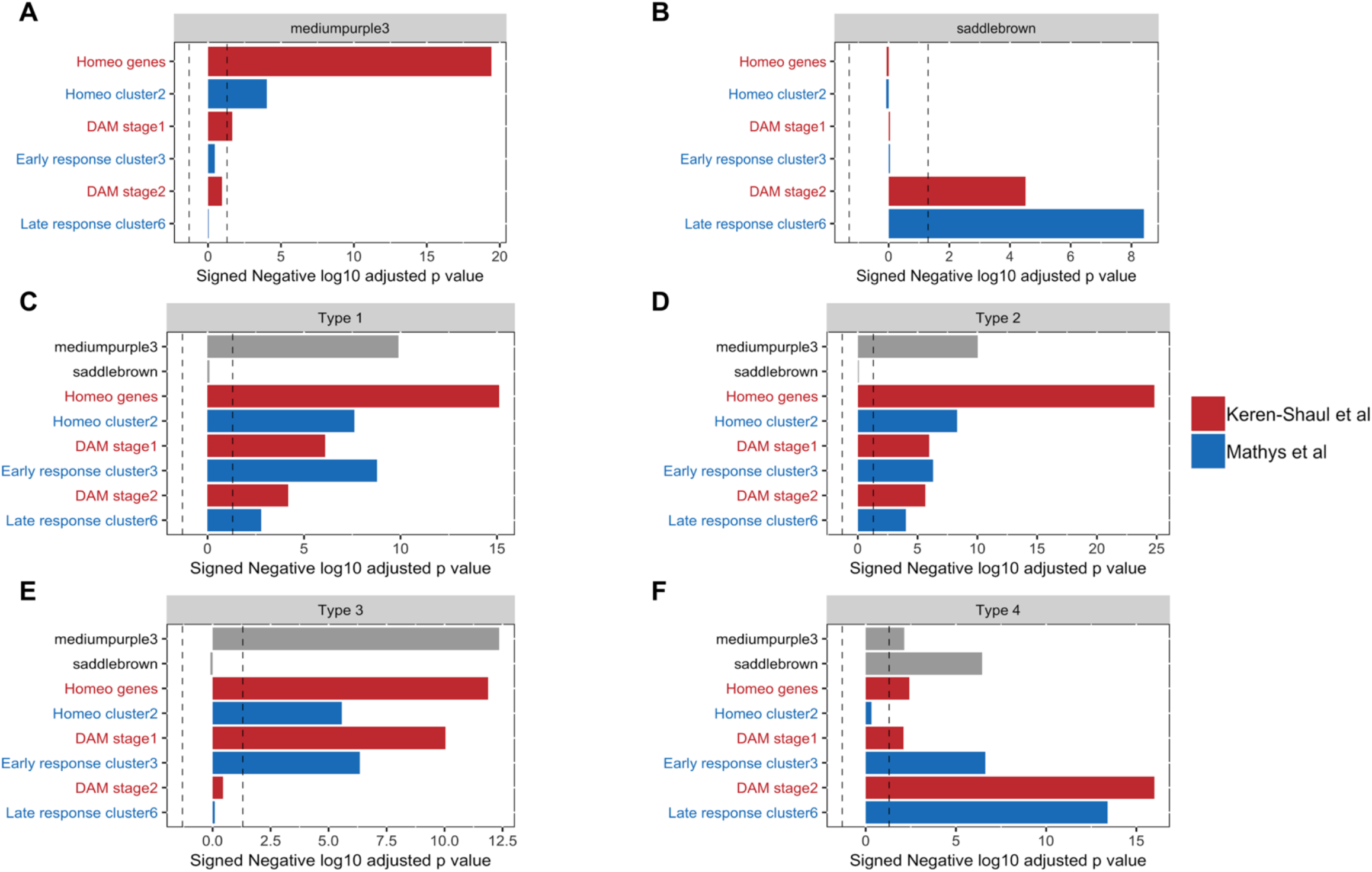
Our microglial modules correlate with known disease-associated microglial gene sets and clusters. **Panels A and B**: The two microglial modules that significantly correlate with AD pathology in our analysis (*saddlebrown* and *mediumpurple3*) also correlate with lists of disease-relevant microglial genes from the mouse model literature (Keren-Shaul et al. ^27^ (red bars) and Mathys et al. ^28^ (blue bars)). The *mediumpurple3* module significantly correlates with homeostatic gene lists from both papers and is more weakly significant with the DAM1 list from Keren-Shaul et al. ^27^. In contrast, *saddlebrown* strongly correlates with the late-stage gene lists from both papers. In addition, *saddlebrown* is itself significantly enriched for late stage/DAM-associated genes, whereas *mediumpurple3* is significantly enriched for homeostatic genes (Figure 5 panels E and G; Supplemental Table 11). **Panels C-F:** Our modules also significantly correlate with lists of subtype specific genes from human single-cell microglial RNA-seq data ^29^ (see Supplemental Table 8 for all correlation values, p-values, and adjusted p-values from this figure). In panels C, D, and E, subtypes 1-3 correlate primarily with homeostatic and early response genes and our own *mediumpurple3* module in our data, while subtype 4 (which is enriched for genes related to amyloid and Lewy body pathology ^29^) correlates primarily with later-stage genes and the *saddlebrown* module (panel F). See text for details; each panel is separately Bonferroni adjusted -dotted line is p-value = 0.05.

**Figure 5:**
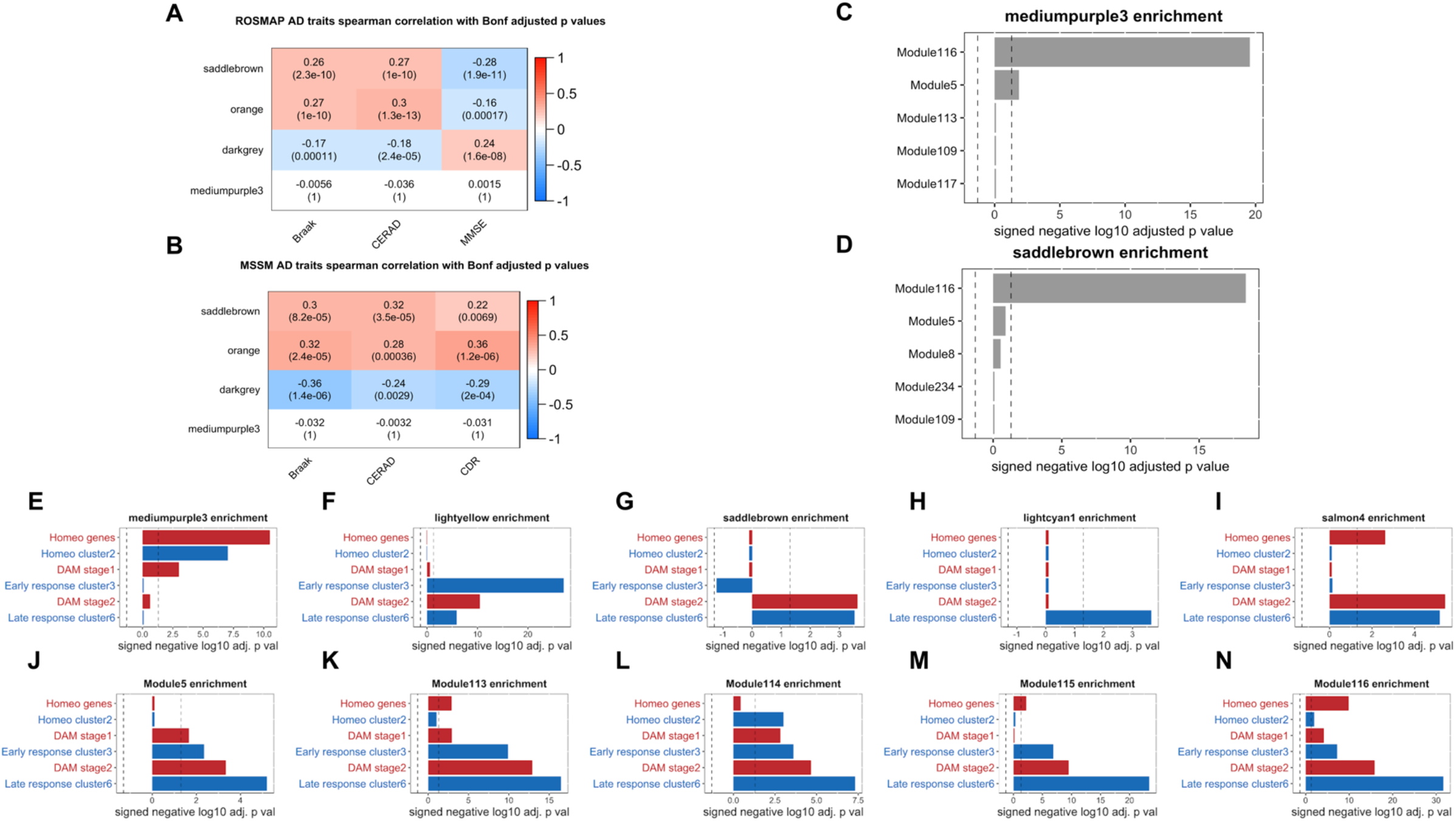
The modules identified in our NPH data also correlate with AD pathology in autopsy cohorts. **Panel A:** *Saddlebrown, orange*, and *darkgrey* correlate with CERAD score, Braak stage, and MMSE score in 596 RNA-seq profiles from the Religious Orders Study and Memory and Aging Project (ROSMAP) Study (representing a range of AD pathologic states) ^1^. **Panel B:** similarly, these modules correlate with CERAD, Braak, and CDR score in RNA-seq profiles from frontal cortex in 183 RNA-seq profiles from the MSSM dataset ^4^. **Panels C and D:** *Saddlebrown* and *mediumpurple3* are both primarily enriched for genes from the same microglial module from Mostafavi et al. ^1^, suggesting that microglial genes in our data are correlating differently than in autopsy cohorts. **Panels E-N:** The distribution of mouse microglial gene lists from Keren-Shaul et al. ^27^ (red bars) and Mathys et al. ^28^ (blue bars) are more segregated in the top five microglial modules from the NPH data (E-I) than in the top five microglial modules from Mostafavi et al. ^1, 31^ (J-N) (see text for details and discussion and Supplemental Tables 9-11 for all correlation and enrichment p-values from these panels). For panels A and B, each row is separately Bonferroni adjusted; for panels, C-N, each panel is separately Bonferroni adjusted - dotted line in panels C-N is p-value = 0.05.

We next compared our modules to recently established cluster types from the human single-cell literature. Specifically, Olah et al. recently published a set of single-cell RNA-seq microglia data from post-mortem and surgically resected human brain tissue FACS sorted for microglia, and identified a set of microglial subtypes through PCA-Louvain clustering ^29^. Subtype defining genes were provided by ^29^ (see Methods), and we determined whether the subtype-defining genes from the microglial groups correlated with either of the microglial modules from this paper. As shown in Figure 4, all three of the “homeostatic” modules identified in Olah et al. ^29^ (subtypes 1, 2, and 3) are correlated with both the *mediumpurple3* module from our work, as well as with homeostatic gene lists from the mouse literature ^27, 28^ (Figure 4, panels C, D, and E). In contrast, subtype 4 (the subtype that is most enriched for genes related to amyloid and Lewy body pathology ^29^), correlates most strongly with the *saddlebrown* module, as well as the late-stage, disease associated gene lists (Figure 4, panel F). While there is a clear trend in this data, it should also be pointed out that the microglial subtypes from Olah et al. significantly correlate with many of the subgroups identified in the mouse literature. Note that although Olah et al did include AD autopsy tissue in this analysis, eight out of the fifteen analyzed samples were epilepsy surgical resections, and future work with larger sample sizes will likely further illuminate the role of different microglia subtypes and microglial transition states in AD and other neurologic diseases. In summary, the data presented in Figure 4 support the hypothesis that our gene expression modules are at least partially tracking an early evolution from homeostatic to late stage/disease-associated microglia in biopsies with early AD pathology.

### NPH modules can be found in other publicly available AD datasets

We next determined how applicable our findings in NPH tissue are for brains with diagnosed AD. The ROSMAP dataset constitutes one of the largest datasets of RNA-seq data from AD neocortex, and so we sought to examine how well our modules from the NPH tissue correlate with pathologic stigmata of AD in this cohort. RNA-seq and associated metadata for ROSMAP ^1^ was downloaded from the AMP-AD Knowledge Portal, and we first regressed out variability in gene expression not correlated with disease-relevant metadata, similar to our processing pipeline for NPH data (see Methods). We first sought to determine whether the modules that correlate with β-amyloid and tau in our NPH data correlate with CERAD, Braak, or MMSE score in the ROSMAP data. As shown in Figure 5 (Panel A), *saddlebrown* and *orange* both have significant positive correlations with CERAD and Braak stage from ROSMAP, while *darkgrey* negatively correlates with these metadata. In addition, *saddlebrown* and *orange* negatively correlate with MMSE and *darkgrey* positively correlates with MMSE in the ROSMAP data, suggesting that these modules may also be related to cognitive decline (see Supplemental Table 9 for correlations and p-values). In summary, our glial modules are positively correlating with pathology and cognitive decline while our neuronal module is correlating in the opposing direction, all of which is consistent with known transcriptional changes in the AD RNA-seq literature ^1, 18, 30^. We also examined an additional dataset of frontal cortex RNA-seq data generated at Mount Sinai ^4^, and found a similar trend to the ROSMAP data (Figure 5, Panel B).

Strikingly absent from both the ROSMAP and MSSM data is any correlation of *mediumpurple3* with AD pathology or cognition. In an effort to further investigate why this module may not correlate with AD pathology in either autopsy dataset, we first examined which of the modules previously identified in the ROSMAP data most overlap with the modules in our NPH data.

Interestingly, both *mediumpurple3* and *saddlebrown* overlap the most with the same module from Mostafavi et al. ^1^; as seen in Figure 5C, this is module 116 (see Supplemental Table 10 for full analysis of the overlap of our four NPH modules of interest with modules from Mostafavi et al. ^1^). Mostafavi et al. identified module 116 as a general microglial module ^1^, and the fact that both our disease-associated microglial module and our homeostatic module overlap primarily with 116 suggests that microglial genes are correlating with one another differently in our data than in ROSMAP.

To further examine this phenomenon, we generated a list of the top five microglial modules from our WGCNA analysis (based on enrichment with human microglial genes from Mathys et al. 2019 ^18^) and compared the relative enrichment of these five modules for different disease-stage gene groups from Keren-Shaul et al. ^27^ and Mathys et al. 2017 ^28^. We then compared this analysis with the distribution of different disease-stage gene groups in the top five microglial modules from the ROSMAP analysis ^31^. As seen in Figure 5, microglial modules from our NPH study largely segregate homeostatic and disease-associated gene groups, with no module showing simultaneous enrichment for homeostatic, early, and late-stage genes (and only one module, *salmon4*, showing both homeosatic and late-stage enrichment). In contrast, the five microglial moduels from Mostafavi et al. are enriched for a far broader range of gene lists identified in the mouse literature, and four out of the five are enriched with a list from every category (i.e. homeostasis, early stage, and late stage; Figure 5, Panels J-N; see Supplemental Table 11 for all enrichment values and p-values). Interestingly, none of the microglial modules from Mostafavi et al. negatively correlate with AD pathology ^1^, in contrast to our homeostatic module *mediumpurple3*. We also examined whether the mouse gene groups themselves correlated with AD pathology in NPH, ROSMAP, and MSSM. As seen in Supplemental Figure 4 and Supplemental Table 12, the homeostatic gene groups defined in mice trend negatively with pathology in NPH data and are either near zero or positively correlate with AD pathology in AD autopsy data. This is again consistent with the finding that homeostatic genes negatively correlate with AD pathology in NPH biopsies and have a more complex relationship with pathology in AD autopsy tissue.

Although the microglial response in mice differs from humans in many important ways, this data suggests that the transition from homeostasis to a disease-associated phenotype is also occurring to some extent in the NPH data, whereas many of these genes are co-correlating in autopsy data. This phenomenon is not only seen in the bulk RNA-seq literature, but is also seen in the single nucleus RNA-seq literature. For example, Zhou et al. (2020) ^30^ have recently directly compared single-nucleus RNA-seq data from a mouse AD model vs. human AD autopsy tissue. The authors replicated the down-regulation of homeostatic genes in AD mice, as well as a lack of this phenomenon in human AD autopsy tissue (indeed, the authors even noted an upregulation of homeostatic genes in human AD autopsy tissue). While the microglial response to AD pathology in humans is clearly different from rodents in many respects, there is a clear trend in our data that previously identified homeostatic genes in rodent models of AD segregate separately from disease-associated genes, and negatively correlate with AD pathology in the *mediumpurple3* module. Although the reason why our data more closely models the mouse AD literature is not completely clear, there are several reasons that could reasonably be considered (see Discussion).

### NPH modules correlate with microglial histologic features

Finally, in an effort to more fully elucidate the relationship between microglia, β-amyloid pathology, and our modules, we performed IBA-1/β-amyloid dual staining in NPH biopsies that we sequenced, and correlated our microglial gene expression modules with microglial morphology in the same biopsies. Microglia in biopsies with β-amyloid plaques tended to be more compact, with an activated, ameboid-like morphology (Figure 6A and 6B), and microglial compactness moderately correlated with plaque area in the same biopsy (r = 0.3; p-value = 0.01). Compactness also correlated exclusively with the *saddlebrown* module after Bonferroni adjustment (Figure 6C; see Supplemental Table 13 for all correlations and p-values). In addition to *mediumpurple3*, we also assessed the correlation of the other three microglial modules from our NPH data shown in Figure 5. Interestingly, even though several other modules are enriched for disease-associated genes identified in the mouse AD literature, only *saddlebrown* significantly correlates with microglial compactness. The correlation of *saddlebrown* with microglial morphology is also seen whether or not samples with AD pathology come from patients that report cognitive symptoms. This suggesting that microglial activation and upregulation of genes in the *saddlebrown* module is an initial response that does not immediately lead to cognitive impairment. Also notable is that *mediumpurple3* is not significant in any comparison, suggesting that loss of homeostatic genes is not related to microglial compactness in these biopsies.

**Figure 6:**
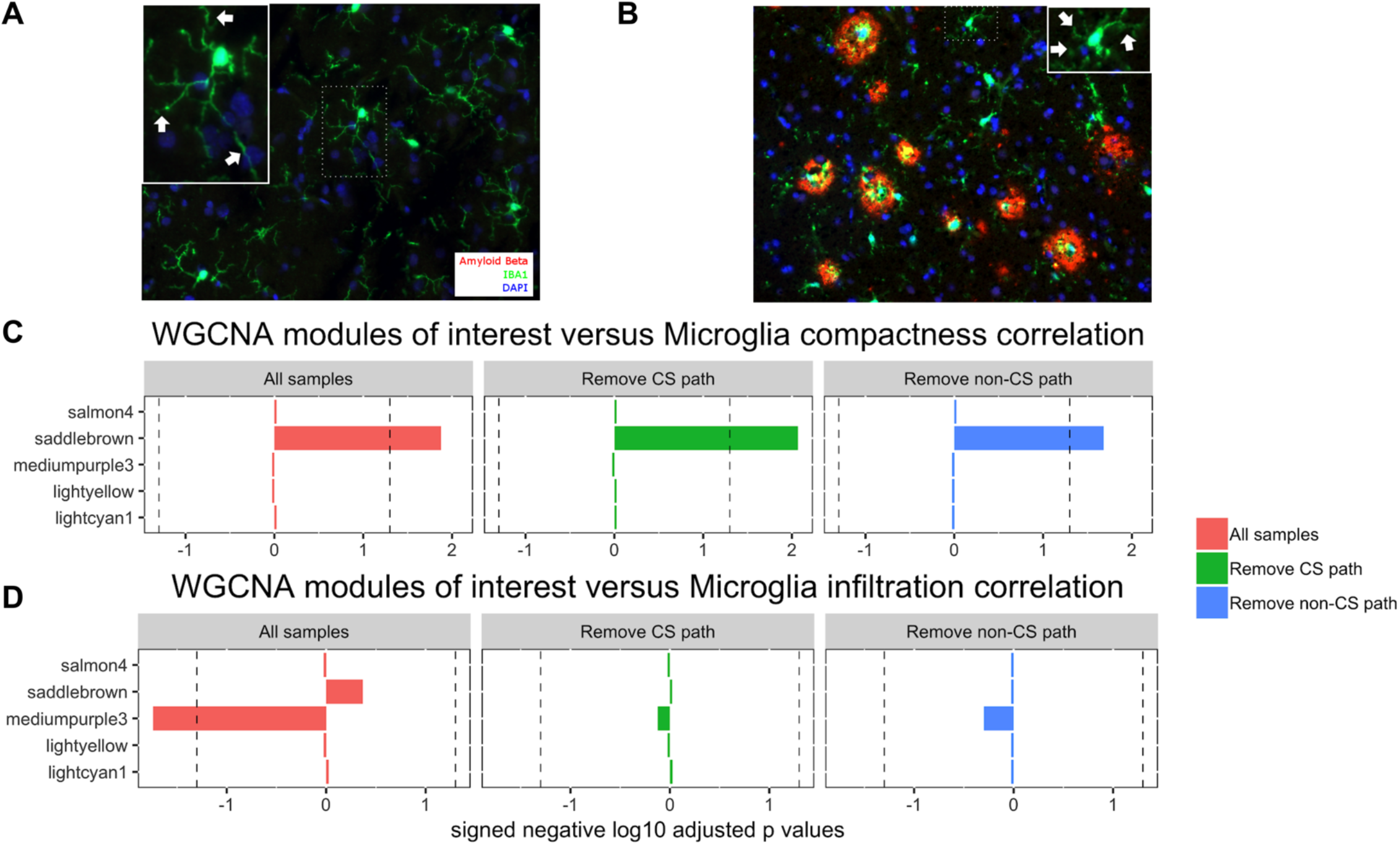
Our microglial modules correlate with microglial morphology and location. We performed β-amyloid/IBA-1 dual staining on selected cases from our cohort. **Panel A**: Cases without β-amyloid showed microglia with longer processes (white arrow; microglial cell in inset is blow-up from area demarcated with dotted line), while (**Panel B**), cases with β-amyloid pathology tended to show more compact, activated microglia. **Panels C and D**: We assessed the correlation of the top five microglial modules shown in Figure 5 panels E-I for microglial compactness (panel C) and microglial β-amyloid plaque infiltration (panel D). Only *saddlebrown* significantly correlates with microglial compactness, and this is seen whether or not samples with AD pathology come from patients that report cognitive symptoms (the green and blue bars represent groups formed using the same methodology as in Figure 3 panels A and B). In addition, only the *mediumpurple3* module significantly (and negatively) correlates with microglial plaque infiltration, suggesting that plaque infiltration correlates with loss of homeostatic genes. When we eliminate samples with AD pathology based on cognitive status we lose significance for *mediumpurple3*, which suggests that this correlation is somewhat weak in this dataset, and also that both sample groups are likely contributing to the overall significance of this finding in the full dataset. In Panel C, n = 59 for all samples (red bars), n = 35 in remove CS path group (green bars), and n = 42 in remove non-CS path group (blue bars); in Panel D, n = 29 for all samples, 11 in remove CS path group (green bars), and n = 18 in remove non-CS path group (blue bars). The three panels in C and the three panels in D are each separately Bonferroni adjusted - dotted line in all panels is p-value = 0.05; see Supplemental Table 13 for all correlations, p-values, and adjusted p-values from this figure.

We also examined the relationship between microglia and plaques in our biopsies, and the relationship with module behavior. Specifically, we looked at the degree of microglial plaque infiltration, and normalized by the total plaque area in each analyzed image (Figure 6, panel D, see Methods for details). Interestingly, the *mediumpurple3* module is the only significant module, and negatively correlates with microglial plaque infiltration, suggesting that plaque infiltration correlates with loss of homeostatic genes. In contrast, the other microglial modules (including *saddlebrown*) show no significant correlation with plaque infiltration. When we eliminate samples with AD pathology from patients based on cognitive status we lose significance for *mediumpurple3*, which suggests that this correlation is somewhat weak in this dataset, and also that both sample groups are likely contributing to the overall significance of this finding in the full dataset. In summary, the associations of *saddlebrown* and *mediumpurple3* with microglial morphology and plaque infiltration do not seem to be sensitive to patient cognitive status, in contrast to the association of these modules with accumulating AD pathology. This is consistent with the view that an initial microglial response to AD pathology is eventually maladaptive, and is associated with accumulating pathology, non-microglial cell responses, and cognitive dysfunction (see Discussion).

## Discussion

In this study, we have used cortical biopsies from hydrocephalus patients to examine changes in gene expression that accompany early-stage AD pathology. The primary findings from this effort are, 1) In contrast to large AD autopsy cohorts, a limited set of mostly microglial genes are found to correlate with AD pathology in our biopsies, 2) A limited set of microglial and non-microglial modules correlate primarily with AD pathology in patients with subjective cognitive impairment, 3) The microglial modules identified in this analysis correlate in a coherent way with homeostatic and disease-associated gene expression lists from the mouse model literature and with microglial subtypes from the human single cell literature, 4) In contrast to the existing AD autopsy literature, these microglial modules replicate the decrease in homeostatic genes in parallel with an increase in late-stage disease-associated genes seen in the mouse AD literature, 5) When analyzed in comparison to IBA1-β-amyloid dual stained sections, our microglial modules correlate with plaque infiltration and microglial morphology, and this change is not sensitive to cognitive status. Taken together, these data suggest that an initial microglial response to AD pathology is eventually maladaptive, and is associated with accumulating pathology, non-microglial cell responses, and cognitive dysfunction.

As noted in the Results section, there is a clear trend in our data that previously identified homeostatic genes in rodent models of AD segregate separately from disease-associated genes, and negatively correlate with AD pathology in the *mediumpurple3* module. Although it is not entirely clear why our data more closely aligns with the mouse AD literature, there are several important differences between our data and human AD autopsy data that may influence the differences we are seeing. One obvious difference is that our tissue was obtained in a different way from AD autopsy tissue. Specifically, our tissue was removed from living cortex while the patient is under anesthesia, as oppposed to being aquired after patient death. In this regard, the lack of any post-mortem artifact in our data makes it more similar to tissue obtained from laboratory mice. Although this presents an intriguing possiblity for how our data may be more similar to the mouse literature, it is difficult to rationalize how post-mortem interval would selectively ablate the decline in microglial homeostatic genes in AD brain tissue, while leaving other changes in gene expression similar between our data and the autopsy data. An alternative, somewhat more plausible explanation is that hypoxia and/or sepsis at the end of life could be affecting the CNS inflammatory response in a way that obscures the AD-associated downregulation of homeostatic genes.

Of course, we should also note that our biopsies represent an earlier stage of pathology than most autopsy cohorts. Note that other studies have included “early AD” groups, although these groups often have more pathology on average than our cohort. For example, Mathys et al. ^18^ has an “early AD” cohort in their study. However, 8 of the 15 subjects in this group are Braak stage 5, which would indicate significant cortical tau pathology in half of their “early AD” subjects (although this is less pathology than in their late-stage subjects). Indeed, Mathys et al. directly compaired their findings to the mouse literature, and the microglial sub-group that was overrepresented in AD had overlap with the late stage/disease associated group. As noted in Figure 5, microglial gene groups from the mouse literature co-correlate in the ROSMAP modules. This raises the possibility that a general microgliosis and increased microglia infiltration may cause the apparent reversal of the decline in homeostatic genes in later-stage AD. However, as already noted earlier, Zhou et al. ^30^ have replicated the down-regulation of homeostatic genes in AD mice, as well as an upregulation of homeostatic genes in human AD autopsy tissue using single-nucleus RNA-seq. This eliminates the possibility that increased homeostatic gene expression in later-stage AD is entirely due to changes in cellular composition, and points to an actual change in microglial gene transcription. Confounding all of these analyses is the additional issue that all of these autopsy cohorts are also older than ours. For example, the mean age of the ROSMAP dataset is 88.7 ^1^, the mean age of the MSSM dataset is 84.7 ^4^, and the Mathys et al. cohort has an average age of 86.7 for the AD group and 87.1 for the non-pathology group ^18^.

Although we cannot resolve all of these issues in this paper, we do think that our data suggests that the downregulation of homeostatic genes in conjunction with an increase in late stage/disease associated genes is occuring at a very early stage, before these patients are actually diagnosed with AD. The implication of this work is that the AD mouse literature may be modeling the earliest stages of the microglial response in AD in a more faithful way than previously recognized. On a more speculative level, one could hypothesize that a later “reversal” and up-regulation of homeostatic genes in the setting of elevated disease-associated genes may be part of the microglial maladaptive response seen in the later stages of AD. If true, how much this correlates with versus causes the later stigmata of disease is an intriguing question that could be explored in future work.

It should be noted that three of the top five microglial modules in our WGCNA analysis by enrichment (*lightyellow, lightcyan1*, and *salmon4*) do not correlate with any of the pathologic metrics in this paper, despite being significantly enriched for several groups of disease-associated genes from the mouse literature (Figure 5). This highlights an obvious point that the microglial response in mice is not exactly the same as in humans. However, it also suggests that the data in this paper can serve as a conceptual bridge between some of the early responses seen in the mouse literature and the human AD literature. While there will always be interspecies differences in these comparisons, the closer similarity of our data to the mouse literature suggests that our data may help clarify which aspects of mouse biology may be accurately modeling the early microglial response in humans. Our work also identifies non-microglial genes as being important in the earliest stages of AD. Although significant genes at the individual level are mostly microglial (Figure 2), WGCNA identifies additional genes that are co-correlating with pathologic changes in these biopsies, and only two out of our four significant modules are “microglial” when we compare these modules to cell-type specific gene lists.

Interestingly, the evolution of homeostatic and disease-associated modules in our data correates with plaque infiltration and an activated morphology, and this change is not sensitive to cognitive status. This suggests that an initial microglial response to AD pathology is eventually maladaptive. Specifically, one could speculate that as the inflammatory response continues and microglial clearance of AD pathology becomes impaired (through a still unknown mechanism), the inflammatory response becomes more robust and additional gene networks (*orange* and *darkgrey*) become significant. These modules also all start to correlate with AD pathology itself as protein clearance is impaired. One might therefore look at the *orange* and *darkgrey* modules for astrocytic and neuronal genes that are involved in early cognitive decline in the setting of a prolonged microglial response. Although speculative (note that we have not proven this sequence of events in this manuscript), this interpretation fits with the prevailing view that while initially playing a positive role in protein clearance ^32, 33^, prolonged neuroinflammation is eventually associated with microglial dysfunction, and this may play a crucial role in mediating cognitive decline in AD ^34, 35, 36^.

Human tissue samples are often aquired for clinical and circumstantial reasons beyond the control of the researcher, and the tissue used in this study also has important limitations for similar reasons. Most obviously, all of the patients in this study have the comorbitidy of hydrocephalus. Since fresh brain tissue cannot be ethically obtained from people purely for research, the comorbidity of hydrocephalus is an unfortunate but completely understandable necessity in order to do the study in this paper. Since “hydrocephalus” is present throughout this cohort, it is not easy to disentangle what effect this might have on gene correlations with AD pathology. However, there are several reasons to hypothesize that the effect of hydrocephalus may be minimal when comparing the data in this study to other gene expression studies on human AD brain tissue. First, AD is rarely “pure” in a pathologic sense. In recent years there has been a growing appreciation that dementia in the elderly is often due to several co-morbid conditions in any individual, as vascular disease, Lewy body dementia, and TDP-43 pathology are frequently seen in AD autopsy cohorts ^37, 38, 39^. Thus, “pure AD” is actually less common among patients with dementia than mixed pathology. Previously reported gene correlations with AD pathology were presumably found in spite of these common confounders, and there is no *a priori* reason to expect hydrocephalus to uniquely affect these correlations more than other common confounders. In addition, it should again be noted that although we have found far fewer changes in gene expression in comparison to previous studies of AD brain tissue, we have also found other changes (i.e. a loss of homeostatic genes) that are not found in AD autopsy cohorts, but are found to some extent in AD animal models. Although one could theorize that findings in this study discordant with the AD autopsy literature are due to hydrocephalus, the alignment of these findings with the AD animal literature strongly suggests that we are observing the same phenotypic change that has been documented multiple times in AD animal models.

In conclusion, this study identifies a restricted set of genes that correlate with early AD cortical pathology and subjective cognitive impairment, and points to future directions for research into how microglia may mediate early cognitive decline. In addition, this work identifies NPH patients with AD pathology as a possible “pre-AD” population that may benefit from early intervention in AD clinical trials. Finally, this study suggests that this patient population may be particularly interesting to study prospectively, and future studies will seek to link these gene expression modules with subsequent cognitive decline or cognitive resilience.

## Methods

### NPH biopsy sample collection and histopathology studies

All human tissue was banked under an IRB approved protocol (IRB-AAAA4666), which allows for the distribution of de-identified tissue samples and clinical data to researchers as “Not Human Subject Research.” NPH was originally defined by ventricular dilation with normal CSF pressure, with a classical clinical triad of imbalance/ataxia; cognitive impairment, particularly short-term memory decline; and urinary incontinence ^40, 41^. The usefulness of the concept of NPH has been more recently challenged ^42, 43^ due to the fact that the symptoms in this clinical triad are common in the elderly population at large ^43^; that (conversely) all three symptoms may not actually occur in all patients diagnosed with NPH ^43, 44^; and that the CSF pressure in NPH patients can actually be variably elevated ^6, 45^, suggesting a chronic deficit in CSF resorption.

Not surprisingly, “NPH” may sometimes arise secondarily to known medical conditions that affect CSF resorption, leading to the concept of “secondary NPH”, to be distinguished from “idiopathic NPH”, or iNPH ^5, 6, 42, 43^. Patients suspected of having NPH can be clinically stratified into “Probable NPH” vs. “Possible NPH”, depending on how well the patient’s symptoms match the appropriate clinical picture and whether any co-morbidities may also be accounting for the patient’s clinical and imaging findings ^44^. We performed RNA-seq on 106 biopsies from NPH patients, 90 of which fell within the definition of “probable” or “possible” iNPH, according to the criteria of Relkin et al. ^44^ (16 biopsies came from patients with a known benign lesion near the cerebral aqueduct that may have contributed to the patient’s chronic hydrocephalus, which would qualify as “secondary NPH”). Our cohort of 106 samples has an average age of 74.9 (standard deviation 8), with 42 females and 64 males. In all cases, patients were shunted for chronic hydrocephalus by the same surgical team, and biopsies were taken from either frontal cortex (middle frontal gyrus at coronal suture) or parietal cortex (∼4 cm off midline in parietal lobe, just above parietooccipital junction). The decision to shunt/biopsy in frontal or parietal cortex was made by the surgeon based on cosmetic considerations; in total, approximately 2/3 of our cohort have frontal biopsies and 1/3 have parietal biopsies. Cortical biopsies were divided in the operating room immediately after removal, and half of each biopsy was frozen in liquid nitrogen while the other half was formalin fixed and paraffin embedded for subsequent pathology diagnosis (Figure 1).

In addition to a hematoxylin and eosin stain, immunohistochemistry was performed with antibodies against tau (AT8; Thermo Fisher; Catalog # MN1020), β-amyloid (6E10; BioLegend; Catalog # 803003), α-synuclein (KM51; Leica; Catalog # NCL-L-ASYN), and TDP-43 (C-terminal rabbit polyclonal; Proteintech; Catalog # 12892-1-AP). All slides were counterstained with hematoxylin. Immunostaining was performed in the Ventana automated slide stainer without manual antigen retrieval and was detected using the Ventana ultraView universal DAB detection kit (Tucson, AZ) as recommended by the manufacturer. Patients in this study had variable amounts of β-amyloid and tau pathology, and no α-synuclein or TDP-43 pathology, or any other visible diagnostic abnormality on hematoxylin and eosin staining. β-amyloid plaques were counted per square mm; in slides with enough tissue, three fields were averaged together, whereas in slides with less tissue, the largest number of possible fields were counted. For tau quantification, we devised a rating scale to grade the minimal degree of tau pathology seen in NPH biopsies (see Supplemental Figure 1). Grade 0 was given to biopsies with no tau pathology. Grade 1 was given to biopsies that have any tau pathology at all, usually one or more dystrophic neurites, but do not make criteria for Grade 2. Grade 2 was given to biopsies that have at least one tau-positive neuron or neuritic plaque, but do not make criteria for Grade 3. Grade 3 was reserved for biopsies with tau pathology evenly distributed throughout the biopsy.

Note that a Braak stage cannot be assigned to these biopsies, as Braak staging is a global assessment of tau pathology that takes into account the presence and density of tau in multiple regions ^46^. Although tau begins to spread broadly into neocortex at Braak stage 4, AD tau pathology shows natural variability from case to case, and occasional focal staining of higher Braak stage areas can be observed at earlier Braak stages ^46^. Thus, in the absence of additional anatomical data Braak staging is not possible. Moreover, it should be emphasized that the primary purpose of the tau grading system devised in this manuscript is to measure the local density of pathology and compare it to local changes in gene expression. All of these considerations apply to Thal staging of β-amyloid as well (i.e. Thal stage requires a global assessment of β-amyloid burden) ^47^. In this case, note that β-amyloid appears initially in neocortex (Thal stage 1), so all of our biopsies with β-amyloid would qualify as Thal stage 1, and we cannot stage higher using only neocortical tissue.

Additional patient data (sex, age, NPH diagnosis, and subjective cognitive status on intake exam) were gathered from the medical record. Cognitive exam data was gathered from patient medical exams as close to the time of biopsy as possible. If possible, an exam note was located where the patient was asked whether they had experienced subjective cognitive impairment. Using this simple metric (yes vs. no), we were able to assign 93 of our sequenced biopsies into “yes” or “no”, with 59 replying “yes” and 34 replying “no” (the remaining 13 biopsies came from patients with no clear answer from the medical record). The average time between exam and biopsy amongst all 93 samples was 120 days.

### NPH sample RNA Sequencing and data preprocessing

RNA was extracted from biopsy samples using miRNeasy Mini Kit (QIAGEN; Cat No./ID: 217004), which purifies total RNA including miRNA (note that our library prep protocol selects poly(A) coding mRNA). RNA integrity was measured on an Agilent Bioanalyzer, and samples with RIN values ≥ 6 were selected for sequencing. RNAs were prepared for sequencing using the Illumina TruSeq mRNA library prep kit, and samples were sequenced with Illumina HiSeq 2000, 2500, and 4000 (potential batch effect attributable to different sequencers was regressed out along with other confounding variables using surrogate variable analysis; see below), and all samples underwent single-end sequencing to 30M read depth. The quality of all fastq files was confirmed with FastQC v 0.11.8 ^48^. Fastq files that passed quality check were further mapped to Genome Reference Consortium Human Build 37 (GRCh37) reference genome with STAR ^49^ (default settings). Output BAM files from STAR were further processed with featureCounts ^50^ to obtain raw counts for each gene of all the sequenced samples.

Genes with less than 5 counts in at least 90% of all samples were first filtered out from the raw count matrix, followed by variance stabilizing Transformation (VST) on filtered counts utilizing the *varianceStabilizingTransformation* function from DESeq2 R package ^51^. Variance stabilizing transformation (VST) is a widely applied strategy that transforms data from a distribution of fluctuating variance into a new distribution with nearly constant variance in order to facilitate downstream analysis ^52^. After VST, surrogate variable analysis (SVA) ^14^ was used to identify variation in gene expression not attributable to β-amyloid load or tau load. Specifically, a full model (with primary variables to be kept) and a null model (with intercept only) were built using the model.matrix() function, and then sva() function from sva r package was applied to determine “surrogate variables (SVs)” from the VST processed gene expression matrix, in which sva function parameter dat was assigned with the gene expression matrix, mod as the full model, mod0 as the null model and method = “irw”. The identified SVs, which represent known and unknown confounding variables in our dataset, were later regressed out with *removeBatchEffect* function from limma R package ^53^. The filtered, VST processed and surrogate variable regressed count matrix was used for all downstream analyses in this manuscript. As an additional exercise, we also attempted to regress out confounding variables individually. To do this, the NPH expression matrix was first quantile-normalized, followed by log2 transformation, and then the batch effect was removed through the ComBat R function from the sva package, and finally age, gender, and RIN were regressed out using the removeBatchEffect R function from the limma package. This method did not yield any significant genes at an FDR threshold of 0.1, confirming the relatively minimal transcriptomic changes in this tissue cohort. This also highlights the need to regress out both known and hidden confounders in our dataset in order to detect the relatively minimal changes in gene expression that we do find.

For our analysis below, we combine RNA-seq data from all biopsies to achieve higher statistical power; this analysis is effectively investigating changes in gene expression shared by two areas of neocortex related to early AD pathology. This power is necessary to detect the relatively subtle changes in gene expression accompanying these early changes in AD pathology (see below). Analyzing frontal and parietal areas separately showed that both analyses trended in a way similar to the combined dataset (see Supplemental Table 1).

### RNA-seq data analysis and module characterization

For single gene analysis, we calculated the Spearman’s correlation between β-amyloid and tau burden and individual genes using the *cor*.*test* function in R. The p-values were further Benjamini-Hochberg (BH) adjusted across all the genes in the dataset using the *p*.*adjust* function in R. To generate gene expression modules, we utilized <u>W</u>eighted <u>G</u>ene <u>C</u>o-expression <u>N</u>etwork <u>A</u>nalysis (WGCNA) to identify gene co-expression modules ^19^, on the default (unsigned) setting, with softPower = 7 and minModuleSize = 20. The eigengene for each WGCNA module was correlated with β-amyloid and tau burden and seven cell-type specific signatures from the mouse literature ^17^ (Neuron, Oligodendrocyte precursor cells, Newly formed oligodendrocytes, Myelinating oligodendrocytes, Astrocytes, Microglia and Endothelial cells), or eight cell-type specific signatures from the human literature ^18^ (Inhibitory neurons, Excitatory neurons, Oligodendrocyte precursor cells, Oligodendrocytes, Astrocytes, Microglia, Endothelial cells and Pericytes). Note that for all correlations of WGCNA eigengenes with other gene lists (including the cell-type specific gene lists above and mouse and human microglial sub-types below), the WGCNA eigengene is correlated with the mean gene expression vector of these various lists. While the eigengene is the preferred way of capturing the variance of highly correlated WGCNA gene modules ^19^, there is no *a priori* reason to assume that the eigengene will capture the majority of the variation in gene lists that are not formed though measuring intercorrelations, and thus, may not be as highly intercorrelated as WGCNA modules. Thus, for cell-type and microglial gene lists we use the mean gene expression vector as a more holistic measure of variation of these gene lists.

For mouse cell-type specific gene signatures, the top 500 differentially expressed genes with average FPKM > 20 and Fold Change (FC) > 1.5 against all other cell types were selected as cell-type specific genes for each cell type. To identify cell class-specific genes from single-nucleus RNA-seq data from the Mathys et al. (2019) study ^18^, we first used edgeR to calculate differential expression among all pairs of “broad cell classes” annotated in the study; each individual nucleus was treated as a “sample” in edgeR, and un-normalized count values were used. For each of these pairs, we then selected cell class-specific genes as those having at least 1.5 positive fold-change between the class of interest and the mean TPM (Transcripts per million) value of all other classes in a pairwise fashion, with the added condition that a given candidate gene mean TPM value should be larger or at least equal to 20. The selected genes were further ranked in descending order based on fold-changes, from which the Top 500 genes were used for correlation analysis. Both mouse and human cell-type specific signatures were calculated as the mean expression of the top 500 cell-type specific genes.

In addition, Fisher’s exact test (FET) was performed to check the enrichment of each mouse and human cell type for the group of individual genes that passed FDR threshold or specific modules of interest. For enrichment analysis, all cell-type specific genes that passed the filtering criteria for FPKM (Fragments per kilobase per million) and FC were selected for the test. The FET p-values were Bonferroni corrected across all cell-types from mice ^17^ or human ^18^ for each comparison. The ontology analysis was performed with all the genes in a given module using over-representation analysis under the “gene set analysis” tab from ConsensusPathDB web tool ^54^. All the level 2 and 3 ontology groups, including all three categories (molecular function (m), biological process (b) and cellular component (c)), with at least two shared genes with the test module were selected and enrichment p-values were calculated through Fisher’s exact test. The q-values were FDR adjusted across all the selected ontology terms within each category (m, b or c). All selected ontology groups were further ranked in ascending order based on q-values, and the top 10 most significantly enriched pathways were listed in Figure 3 (E-H).

Note that for this study, we have employed an FDR threshold of 0.1 when screening large numbers of hypotheses to identify genes or modules of interest (i.e. genes or WGCNA modules that correlate with AD pathology or ontology groups that are enriched for module genes). Throughout the rest of the manuscript, tests of hypotheses about these genes/modules have employed a Bonferroni correction with a p-value threshold of 0.05.

### Comparison of NPH modules with AD autopsy datasets

We compared NPH datasets with two publicly available AD autopsy datasets:

1) The Religious Orders Study and Memory and Aging Project (ROSMAP) Study (Synapse ID: syn3219045):

RNA-Seq raw counts of in total 596 human subjects were obtained from paired-end fastq files using STAR + featureCounts pipeline for pair-ended reads. The raw counts matrix was filtered and VST transformed as described above. Finally, variation in gene expression not attributable to Braak stage, CERAD score, and MMSE was regressed out using SVA, similar to our processing pipeline for NPH data.

2) The Mount Sinai Brain Bank (MSBB) Study (Synapse ID: syn3159438):

We downloaded 183 bulk RNA-Seq raw counts tables from human brain samples from Brodmann (BM) area 10. Samples were preprocessed the same way as our NPH samples and the ROSMAP data, and SVA was used to regress out variability not attributable to Braak stage, CERAD score, and CDR score.

### Effects of cognitive status (CS) on NPH AD traits

In Supplemental Figure 2, patients that report subjective cognitive impairment are compared to patients that report no cognitive impairment. The cognitive information of each subject was obtained as described above. Mann-Whitney U test was first performed to examine the difference in average β-amyloid and tau load between subjects who report cognitive impairment and subjects who report no cognitive impairment. To further explore the distribution of these variables, we first performed a Fisher exact test on whether tau and β-amyloid significantly co-occur in biopsies from patients with reported cognitive impairment vs. biopsies from patients that report no cognitive impairment. We next performed a Mantel-Haenszel test of whether the odds ratios between the two groups were different (Supplemental Figure 2). In Figure 3 panel A and B, we asked whether samples with AD pathology from either group were significantly driving the correlation between the modules and AD pathology. To do this, we first removed all samples with AD pathology from our cohort where patients reported cognitive impairment (our “Remove CS path” group, n = 66). Next, we removed all samples with AD pathology where patients that reported no cognitive impairment (our “Remove non-CS path” group, n = 80). Note that in both groups, all biopsies without AD pathology were included. Finally, we ran 1000 iterations where half of the samples with AD pathology from “Remove non-CS path” group (20 out of 40) were randomly selected to be replaced by another randomly selected 20 samples with pathology from “Remove CS path” group to form an artificial “Remove non-CS path” group (i.e. pathology samples with subjective cognitive impairment are being randomly replaced with pathology samples without documented cognitive impairment). Mann-Whitney test of β-amyloid or tau between the real and artificial groups and correlations of module eigengenes with β-amyloid or tau in both groups were performed for each iteration and p-values from these tests were recorded. The correlation p-values were further BH adjusted across all WGCNA modules. At the end of all iterations, the number of times out of 1000 iterations 1) the Mann-Whitney test p-value was less than 0.05 for β-amyloid or tau and 2) the BH adjusted correlation p-values that passed significance threshold (0.1) for any of the four modules in Figure 3 was reported. As noted in Supplemental Table 4, this did not statistically change the overall burden of pathology in any of the simulations. In contrast, all four modules of interest fail to pass 0.1 FDR significance in their correlation with β-amyloid and tau for the majority of the simulations.

### Microglia activation stage and subtype specific genes identification

Microglia activation stage specific genes from mouse studies were obtained from two separate datasets. We used three sets of genes reported in Keren-Shaul, et al. ^27^ (Homeostatic genes, DAM stage1 and DAM stage2), and three sets of genes reported in Mathys, et al. ^28^ (Homeostatic cluster 2, Early response cluster 3 (the primary early response cluster), and Late response cluster 6). We defined cell-type specific genes using comparisons done in these manuscripts. Specifically, homeostatic genes from Keren-Shaul et al. were defined as differentially expressed genes with a minimum 2-fold increase in expression compared to Tg 5xFAD samples and FDR less than 0.1. The DAM stage 1 and DAM stage 2 genes were filtered through similar criteria except that the DE genes were calculated based on the comparison between these two DAM stages. For gene lists from Mathys, et al. (i.e. clusters 2, 3, and 6), cluster-specific genes were defined as genes upregulated in a given cluster in comparison to either or both of the other two clusters, and not downregulated in either of these two comparisons. The significance of these comparisons was defined as absolute value of the corrected z score > 1.25, which is equivalent to an FDR corrected p value >0.1. Mouse gene symbols were converted to human gene symbols with R biomaRt package for comparison with the data in this paper.

Microglia subtype information was obtained from a human microglial dataset kindly provided by Olah M, et al. ^29^ and consisted of single cell RNA-Seq results of 14 different microglial cell types obtained from autopsy human cortical brain tissues. A microglia subtype specific gene is defined as having a significant upregulation in the given cell type compared to at least one of the rest of the cell types while no downregulation in the given cell type compared to any of the other cell types.

### Immunofluorescence staining and imaging

Immunofluorescence was performed on cortical biopsies for β-amyloid and IBA-1 to visualize microglial morphology and microglial plaque infiltration. Fixed, paraffin embedded tissue was sectioned at 7 µm and slides were submerged in two washes of Histo-clear II (National Diagnostics HS-202) for 10 minutes each, before being washed in the following series for 1 minute each: 100% ethanol, 100% ethanol, 70% ethanol, and MQ water. They were then washed 3 times in TBS for 5 minutes each. Antigen retrieval was performed by incubating the slides in citrate buffer for 25 minutes in a microwave set to 400 Watts. The slides in citrate buffer were rested at room temperature for 30 minutes before a 5-minute wash with MQ water, two 5-minute permeabilization washes in TBS with 0.25% Triton X-100 (ACROS Organics), and a 5-minute wash in TBS. The sections were blocked with 10% donkey serum (Abcam ab7475) and 1% BSA (Sigma-Aldrich A7284) in TBS for 1 hour at room temperature. A primary antibody solution was created in 1% BSA in TBS, with the addition of mouse polyclonal anti-β-amyloid (1:200; Cell signaling Catalog # 15126, lot 1) and rabbit polyclonal anti-IBA1 (1:500; Wako Catalog # 019-19741, lot ptr2404). The sections were stained overnight in the primary antibody solution at 4°C, after which they underwent three 5-minute washes in TBS. A secondary antibody solution was created in 1% BSA in TBS, with the addition of Alexa Fluor 555 conjugated Donkey Anti-Mouse (1:1000; Invitrogen A-31570, lot 1850121) and Alexa Fluor 488 conjugated Donkey Anti-Rabbit (1:1000; Invitrogen A-21206, lot 1981155). The sections were incubated for 1 hour at room temperature in the secondary antibody solution, after which they underwent three 5-minute washes in TBS. To block autofluorescence, Sudan Black B (EMS 21610) solution at 0.1% m/v in 30% MQ water and 70% ethanol was placed on the sections for 20 minutes, after which they underwent three 5-minute washes in TBS. Nuclei were labeled with DAPI (1:4000 from stock; Invitrogen D1306) in TBS for 5 minutes, before three 1-minute washes in TBS. Slides were mounted with Vectashield (Vector H-1000).

Cellprofiler was used to perform image analysis on the immunohistochemistry stained slides ^55^. IBA1 staining area to determine microglia density was identified with thresholding based on the background variance. Individual Iba1-stained microglial cells were identified with thresholding after successive edge-preserving smoothing to remove stray processes, and the microglia cell shape traced by enhancing neurite shapes and thresholding the resulting image. Plaques were identified with thresholding based on the background variance. Microglial infiltration of plaques was determined from the total Iba1 staining area overlapping plaques for a sample, normalized by total plaque area. Microglial compactness was calculated on individual Iba1-stained cells as a measurement output defined by Cellprofiler as “the mean squared distance of the object’s pixels from the centroid divided by the area.” Calculated this way, this metric is lower for more compact cells; in Figure 6, we invert this value so that our modules positively correlate with “compactness.”

## Acknowledgements

This work was supported by NIH grants K08-AG049938 (AFT) and K76-AG054868 (AFT). We would also like to acknowledge Richard Hickman for assistance with developing the tau grading system.

## Author Contributions

WH, XF, SL, DB, ES, VM, and ZT all contributed to organizing and analyzing the data in this manuscript. AMB and HX assisted with completion and analysis of immunofluorescence experiments. GMK recruited the patients, evaluated them, operated on them, and made sure the tissue was acquired properly and in a timely fashion, and was assisted by GI and RAM. AFT designed the study and subsequent analyses, oversaw its completion, and wrote the manuscript together with WH.

---

**Supplemental Figure 1.**
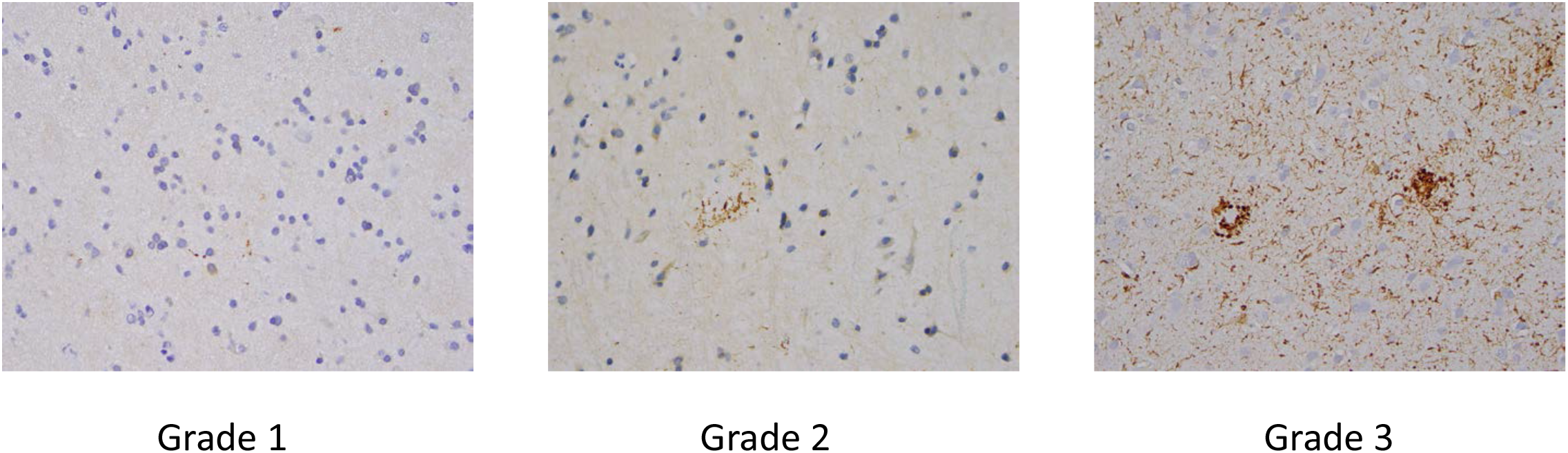
We devised a rating scale to grade the minimal degree of tau pathology seen in NPH biopsies. Grade 0 was given to biopsies with no tau pathology. Grade 1 (left panel) was given to biopsies that have any tau pathology at all, usually one or more dystrophic neurites, but do not make criteria for Grade 2 (middle panel). Grade 2 was given to biopsies that have at least one tau-positive neuron or neuritic plaque, but do not make criteria for Grade 3 (right panel). Grade 3 was reserved for biopsies with tau pathology evenly distributed throughout the biopsy.

**Supplemental Figure 2.**
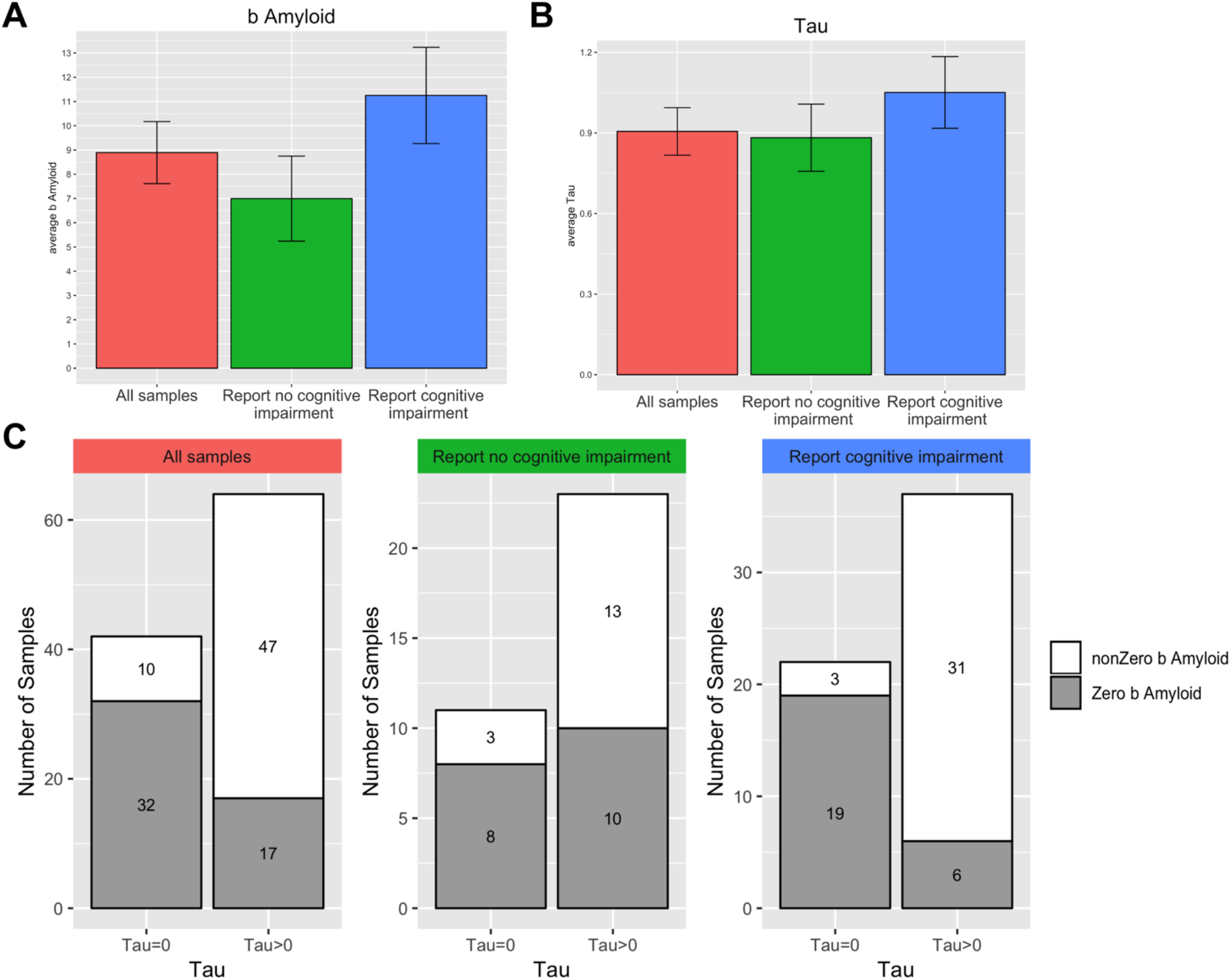
Cognitive status minimally influences AD pathologic burden in this cohort. Panels A and B: Patients who report cognitive impairment have a non-significant trend for higher levels of β-amyloid and tau on biopsy when compared to patients who report no cognitive impairment (error bars = standard error; p-value for β-amyloid = 0.21; p-value for tau = 0.66 by Mann Whitney U test). Panel C: Despite a non-significant difference in overall AD pathology, patients who report cognitive impairment are more likely to have β-amyloid and tau in the same biopsy (Fisher exact test for presence of both β-amyloid and tau in biopsies from patients that report cognitive impairment p-value = 0.0001269, for patients that report no cognitive impairment p-value = 0.2587; Mantel Haenszel test for whether odds ratios between the two groups are different: two-sided p-value = 0.0002021). Note that the presence of 9 samples with no cognitive information and zero tau score in the full cohort make the overall average tau value in panel B (“all samples”) lower than the average value between patients that report cognitive impairment and patients that report no cognitive impairment.

**Supplemental Figure 3.**
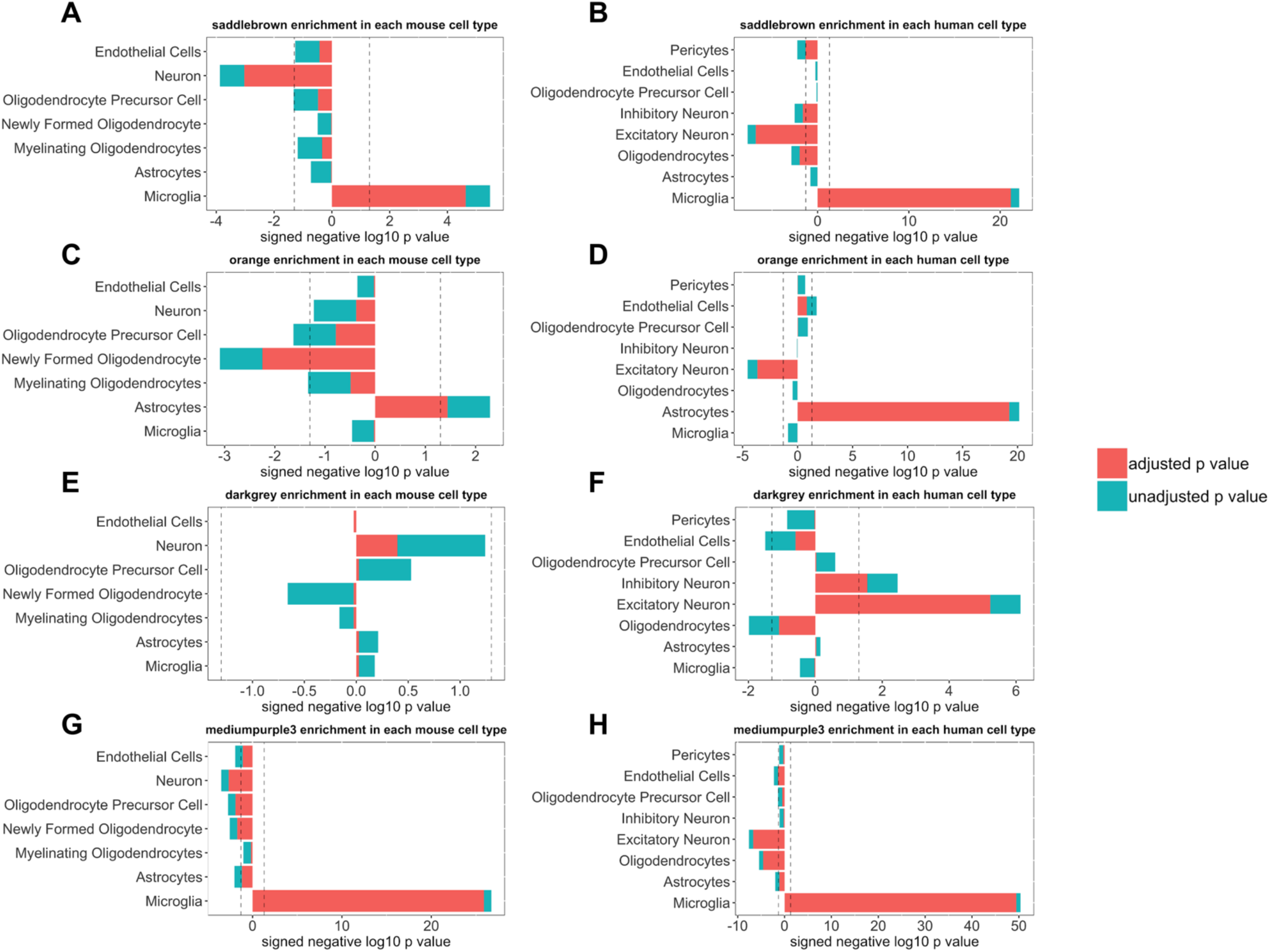
Shown is the enrichment for each of the four modules in Figure 3 with seven cell-type specific signatures from the mouse literature ^17^ (panels A, C, E, and G), or eight cell-type specific signatures from the human literature ^18^ (panels B, D, F, and H) using the Fisher’s exact test; see Supplemental Tables 5 and 6 for full correlation and enrichment analysis of these modules with all mouse and human cell types. Each panel is separately Bonferroni adjusted – both adjusted and unadjusted p-values are shown; dotted line in all panels is p-value = 0.05.

**Supplemental Figure 4.**
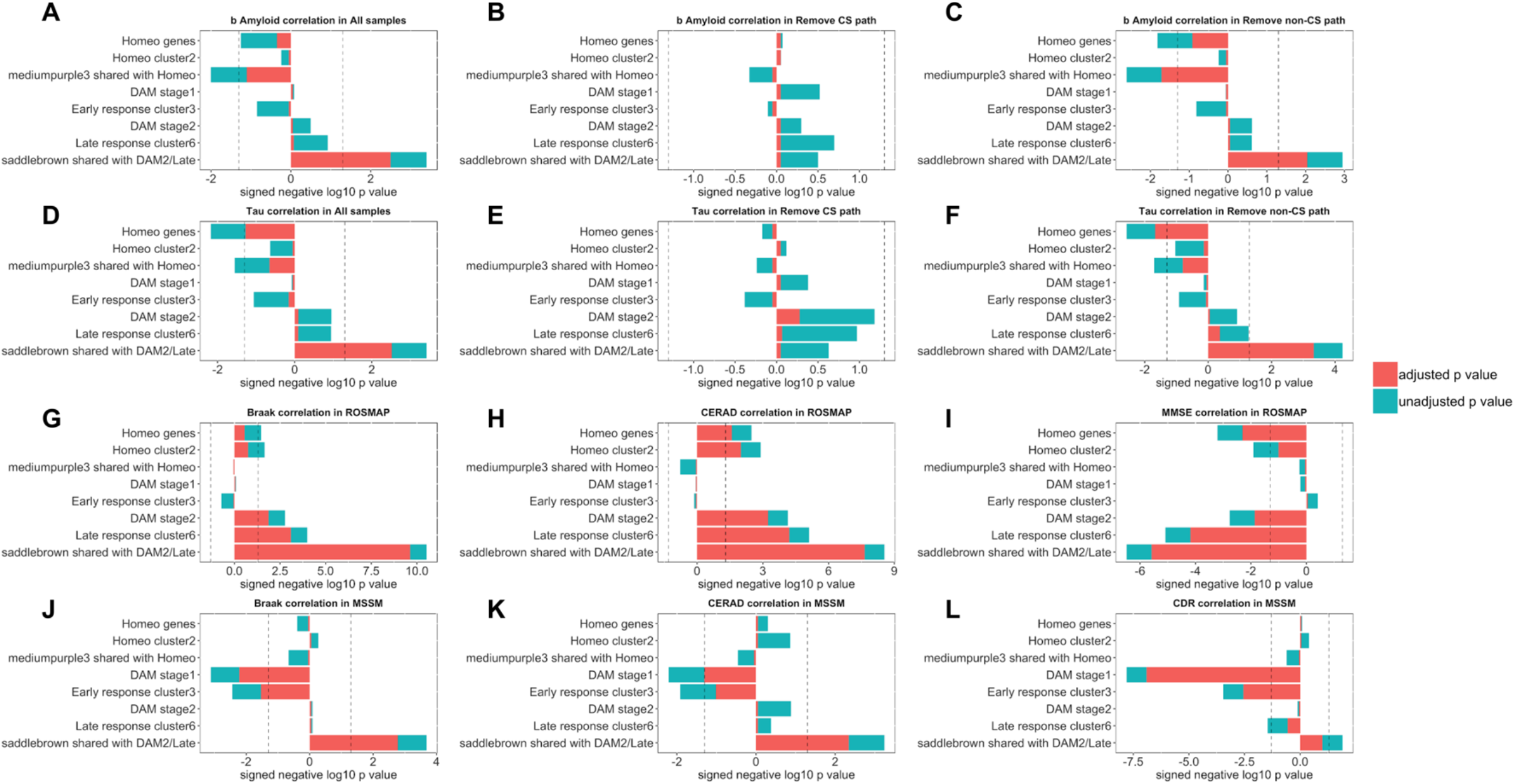
Mouse microglial genes variably correlate with AD pathology in NPH and autopsy tissue. Shown are correlations of three sets of genes reported in Keren-Shaul, et al. ^27^ (Homeostatic genes, DAM stage1 and DAM stage2), and three sets of genes reported in Mathys, et al. ^28^ (Homeostatic cluster 2, Early response cluster 3 (the primary early response cluster), and Late response cluster 6) with β-amyloid and tau in NPH samples (panels A and D) and our remove CS path (panels B and E) and remove non-CS path (panels C and F) groups from Figure 3. Correlation of these mouse modules with AD traits in ROSMAP (panels G, H, and I) and MSSM (panels J, K, and L) are also shown. In each panel is also shown the correlation of the subset of genes in *mediumpurple3* that overlap with the homeostatic lists from Keren-Shaul, et al. ^27^ or Mathys, et al. ^28^, and the correlation of the subset of genes in *saddlebrown* that overlap with DAM stage 2 or late response cluster 6 from Keren-Shaul, et al. ^27^ and Mathys, et al. ^28^ (see Supplemental Table 14 for overlap of these modules with lists from Keren-Shaul, et al. ^27^ and Mathys, et al. ^28^). Note that in NPH data, homeostatic genes negatively correlate with β-amyloid and tau, while late stage/disease associated gene lists from mice positively correlate (although not significantly) with β-amyloid and tau. Not surprisingly, homeostatic and late stage/DAM stage 2 genes that overlap with *mediumpurple3* and *saddlebrown* respectively correlate stronger with AD pathology in the NPH biopsies. Autopsy datasets offer a more complex figure, with ROSMAP data showing positive correlations of both homeostatic and late stage/disease associated genes with AD traits, similarly to what is reported in Zhou et al. ^30^. The MSSM dataset offers a somewhat different picture, with early response and DAM stage 1 genes negatively correlating with AD pathology, while homeostatic and late stage/DAM2 genes show little correlation. Note, however, that although the late stage/DAM2 lists do not correlate with pathology as a whole in the MSSM dataset, the subset of these genes that overlap with *saddlebrown* do strongly correlate with AD traits in the MSSM dataset, consistent with an ongoing microglial response that may not completely correspond to the mouse literature. Each panel is separately Bonferroni adjusted – both adjusted and unadjusted p-values are shown; dotted line in all panels is p-value = 0.05. See Supplemental Table 13 for all of the correlation values, p-values, and adjusted p-values in this figure.

**Supplemental Table 1** -list of all genes that correlate with β-amyloid and tau in the full dataset, also correlations with frontal samples only and parietal samples only, along with p-values and adjusted p-values using the Benjamini-Hochberg procedure. For the top 100 genes that correlate with β-amyloid in the full dataset, the correlation coefficient of these genes with β-amyloid in frontal and parietal samples are themselves highly correlated, with a Spearman’s correlation of r=0.8691309, p<2.2E-16 across all 100 genes. Both frontal and parietal samples are equally similar to the full dataset as well, with a Spearman’s correlation of r = 0.9893544, p<2.2E-16 when the correlation of these 100 genes with β-amyloid are compared between the full dataset vs. frontal samples, and r=0.9297355, p<2.2E-16 when the comparison is between the full dataset and parietal samples. A similar relationship holds with tau; for the top 100 genes that correlate with tau in the full dataset, the correlation coefficient of these genes with tau in frontal and parietal samples have a Spearman’s correlation of r=0.7948915, p<2.2E-16. When the correlation of these 100 genes with tau are compared between the full dataset vs. frontal samples, r = 0.9769622, p<2.2E-16, and when the correlation of these 100 genes with tau are compared between the full dataset vs. parietal samples, r=0.9037152, p<2.2E-16. Also included in this table file are the p-values and adjusted p-values that correspond to Figure 2 panels C and D (Fisher’s exact test enrichment for cell-type specific genes among the genes that individually past FDR 0.1 threshold in the full dataset, both using mouse cell-type specific RNA-seq data ^17^ and human single-nucleus RNA-seq data ^18^).

**Supplemental Table 2** – Correlation of all WGCNA modules with β-amyloid and tau pathology in all samples, in the remove CS-path group, and in the remove non-CS path group from Figure 3, along with p-values and adjusted p-values using the Benjamini-Hochberg procedure.

**Supplemental Table 3** – shown are all genes in all WGCNA modules from this manuscript, along with correlations of module genes with modules PC1 eigengene vectors. In addition, the genes in the four modules of interest from this paper are also shown in separate tabs for easier viewing (i.e. *saddlebrown, orange, darkgrey*, and *mediumpurple3*).

**Supplemental Table 4** – In an effort to further investigate the importance of cognitive status in our samples, we ran 1000 iterations where half of the samples with pathology and reported cognitive impairment are being replaced with samples with pathology and no reported cognitive impairment (i.e. the blue analysis in Figure 3, panels A and B is having half of its pathology samples replaced with pathology samples from the green analysis). As shown in Supplemental Table 4, this did not statistically change the overall burden of pathology in any of the simulations using the Mann-Whitney U test. In contrast, all four of our modules fail to clear 0.1 FDR significance using the Benjamini-Hochberg procedure in their correlation with β-amyloid and tau for the majority of the simulations. Taken together, these findings indicate that the correlations of these modules with AD pathology are highly sensitive to cognitive status.

**Supplemental Table 5** -The eigengene of the *saddlebrown, orange, darkgrey*, and *mediumpurple3* modules was correlated with seven cell-type specific signatures from the mouse literature ^17^ (Neuron, Oligodendrocyte precursor cells (OPCs), Newly formed oligodendrocytes, Myelinating oligodendrocytes, Astrocytes, Microglia and Endothelial cells). In addition, the enrichment for each module for these gene lists using the Fisher’s exact test is shown. For both correlations and enrichment statistics, p-values and Bonferroni adjusted p-values are shown.

**Supplemental Table 6** - The eigengene of the *saddlebrown, orange, darkgrey*, and *mediumpurple3* modules was correlated with eight cell-type specific signatures from the human literature ^18^ (Inhibitory neurons, Excitatory neurons, Oligodendrocyte precursor cells (OPCs), Oligodendrocytes, Astrocytes, Microglia, Endothelial cells and Pericytes). In addition, the enrichment for each module for these gene lists using the Fisher’s exact test is shown. For both correlations and enrichment statistics, p-values and Bonferroni adjusted p-values are shown.

**Supplemental Table 7** – Shown is the full ontology analysis for *saddlebrown, orange, darkgrey*, and *mediumpurple3* modules. In addition are shown the ontology analysis for genes that positively and negatively correlate with the PC1 eigengene for these four modules. Three categories of ontology analysis were investigated (molecular function(m), biological process (b) and cellular component(c)), with all categories included that have at least two shared genes with the test module. The q values were FDR adjusted across all the selected ontology terms within each category(m, b or c); see Methods for details.

**Supplemental Table 8** – Correlations, p-values, and Bonferroni adjusted p-values for all panels in Figure 4 are shown.

**Supplemental Table 9** – Correlations, p-values, and Bonferroni adjusted p-values for panels A and B in Figure 5 are shown.

**Supplemental Table 10** - Correlation of the *saddlebrown, orange, darkgrey*, and *mediumpurple3* modules with all of the modules from Mostafavi et al. ^1^, as well as overlap (enrichment) using the Fisher’s exact test. For both correlations and enrichment statistics, p-values and Bonferroni adjusted p-values are shown.

**Supplemental Table 11** – Correlations, p-values, and Bonferroni adjusted p-values for panels E-N in Figure 5 are shown.

**Supplemental Table 12** – Correlations, p-values, and Bonferroni adjusted p-values for Supplemental Figure 4 are shown.

**Supplemental Table 13** – Correlations, p-values, and Bonferroni adjusted p-values for Figure 6 are shown.

**Supplemental Table 14** – Shown are the gene members from the top 5 microglial modules from our WGCNA analysis, along with correlations with module PC1 eigengenes and membership in microglial gene lists from Keren-Shaul, et al. ^27^ and Mathys, et al. ^28^ (0 = not a member, 1 = is a member).

